# Atmospheric pollution contributed to destabilizing a small mammal community amid four decades of environmental changes

**DOI:** 10.64898/2026.11.03.764082

**Authors:** Thibaut Rota, Oldrich Rakovec, Richard Ševčík, Jan Zima, Vladimír Bejček, Karel Šťastný, František Sedláček, Jiří Šindelář, Adéla Šípková, Vladislav Chrastný, Wesley M. Hochachka, Markéta Zárybnická

## Abstract

How multiple environmental changes affect ecological stability is a pressing question. We analyzed 39 years (1986–2024) of population abundances for six small-mammal species in Europe’s historically air-polluted Black Triangle (Czechia), alongside monthly air pollution (positively correlated with lead (Pb) deposition in tree rings), vegetation greenness (NDVI), climate, and soil moisture data. We used Empirical Dynamic Modeling to identify instability (chaos), species-species and species-environment causalities, time-varying species interactions and feedback loops, and compare out-of-sample forecasts. Late-1990s air cleanup likely triggered a delayed critical transition in the community, showing that post-disturbance recovery can occur as an abrupt state shift rather than a gradual return. Pollution primarily affected dominant omnivores, destabilizing the community by strengthening species interactions but weakening within-species competition. Seasonal vegetation and soil-moisture cycles promoted destabilizing, self-reinforcing feedback loops. Incorporating key environmental predictors improved forecasts of chaotic dynamics, demonstrating that identifying primary drivers is essential to anticipate unstable ecosystems’ futures.

## Introduction

Chaos, defined as the sensitivity of nonlinear systems to initial conditions —whereby a perturbation produces exponentially-diverging outcomes (1)— relates to instability (1, 2), abrupt changes in ecosystem state (3), and a lack of forecasting ability (4, 5), all relevant issues with global change (2, 6). After a longstanding debate on the supposed rarity of chaos (7, 1, 8), modern analysis has concluded that chaos occurred in ∼1/3 of available population time series, especially in *r*-selected species often sensitive to their abiotic environments (8, 9). The focus has now shifted from the mere detection of chaos (8, 10, 11) to assessing how it occurs and what its implications are in a fast- changing world (6, 2, 12, 9).

Dynamical instability, or chaos, is theoretically common in highly-dimensional systems (1), and populations exhibiting high growth rates (7) and strong nonlinear delayed responses to perturbations (13, 14). Network topology also matters (6), such as intransitive feedback loops with their cycles of winnerless dominance (e.g., “rock/paper/scissors”). As shown in a coastal community (6), these networks can fluctuate at the edge of chaos, which is amplified by seasonal forcing. Multiple stressors could cause deterministic chaos and/or critical transitions in altering the physiology, behavior and vital rates of individual organisms, ultimately altering demographics (15, 4, 5). Various facets of a system can be altered by environmental changes: population dynamics (16, 17), time varying-interactions (2, 18), dynamical instability (19, 6, 8) and multi-stability (20–23). Such complexity is hardly amenable to a reductionist approach. Instead, long term studies of entire natural communities and their environments are required (18, 21), with environmental changes and internal states of a system being used to infer the cause of observed dynamics.

Atmospheric pollution, which peaked in the 1960s–1990s in Europe, is associated with toxic metal(loid)s such as lead (Pb), one of the most potent toxic elements on Earth (27). Massive amounts of lead have been emitted from coal burning and leaded gasoline (tetraethyl-lead) before the 1990/2000s bans of the latter (27). In fact, changes in climate, land use, vegetation, and pollution occurred concomitantly in Europe in the last decades, with for instance atmospheric pollution and acidic rains causing forest diebacks in Central Europe (28), amid the onset of climate change (24, 25). The Black Triangle region (at the intersection of Germany, Poland and Czechia) was particularly impacted by brown-coal burning during the late Czechoslovakia-era (28). This induced large diebacks of the Norway spruces, consequent clearcuts, and replanting with Colorado spruce in the late 1980s. Following the Velvet revolution in 1989, air quality improved considerably (31, 32). However, the effect of pollution stayed overlooked in ecology (16), and its potential effects remained entangled with other changes (e.g., climate). This conundrum is blurring our attribution of ecological changes (34, 5), impeding our understanding of the mechanisms driving past, current and future ecosystems.

Small mammals are key to terrestrial ecosystems (35, 36). As *r*-selected species, they exhibit various dynamics: limit cycles, quasi-cycles, chaos (37) and critical transitions whose determinants remain debated (38, 39). While deterministic chaos is hindering mid- and long-term forecast ability (14), identifying its drivers would help reconstruct strange attractors and improve forecasts (15, 34, 1, 5). Considering the outbreaks of field voles damaging saplings and crops (40), and the role of small mammals for transferring energy to higher trophic levels, this issue has implications for management (5). So far, early-forest succession and afforestation —or unknown shared driver(s)— were proposed as causing the dampening of the large-amplitude cycles of field voles in Europe (38, 41), and concomitant increases of yellow-necked mouse and bank voles (41, 36, 39). However, these suggestions overlook the potential role of changing levels of atmospheric pollution and that the last two aforementioned species have been suggested as particularly sensitive to metallic contamination (42, 43).

Here, we analyze 39-years (1986–2024) of semiannual abundance data from a small-mammal community from the Ore Mountains (Black Triangle, Czechia). The network is composed of six main species from three guilds: omnivores (*Apodemus flavicollis,* the yellow-necked mouse, and *Clethrionomys glareolus,* the bank vole), strict insectivores (*Sorex araneus* and *Sorex minutus,* the common and pigmy shrews), and strict herbivores (*Microtus agrestis* and *Microtus arvalis,* the field and common voles). We added to this dataset monthly data on atmospheric pollution (SPM), meteorology, topsoil moisture, and remote sensing vegetation index (NDVI). We evaluated the prevalence of chaos with Lyapunov exponents, and the eventuality of a critical transition using a multiple change-point Bayesian model and multi-index early warning signals (EWS). Using Empirical Dynamic Modeling (EDM) (44), we then tested for causal links between environmental drivers and population dynamics, tested how the environment may affect interactions in the network and lead to chaos. In addition to changes in climate and vegetation, we expected that changes in SPM (SI Fig. S1A) would especially alter community dynamics, since the exposure to Pb differed among guilds (SI Fig. S1B). Hence, we hypothesized that SPM would negatively impact the pollution-sensitive omnivores (42, 43) and the insectivorous shrews showing high Pb burdens (SI Fig. S1B), releasing competition on the less contaminated herbivores (SI Fig. S1B). Thus, the polluted period could be dominated by negative effects of pollution on the omnivores and insectivores, and overcompensatory feedbacks by herbivores in a system less stabilized by omnivory (45, 46). We expected this context to be prone to strengthened interactions among species, positive feedback loops, chaos, and bifurcations (2, 3, 23, 47). Therefore, the recovery of the community after the perturbation could have taken the form of an abrupt community shift. Finally, we hypothesized an improvement of mid-term forecast abilities when adding relevant environmental drivers in our models, indicating a better reconstruction of the chaotic attractor (34).

## Results

### Time series of environmental drivers

Atmospheric pollution (SPM) was high from 1985 to 1998, then dropped by ∼80% in the 2000s, remaining low until now (Fig. 1). During the polluted period, the recurrence frequency of pollution pulses ranged from 8 to 30 months (Fig. 1A). SPM had the strongest temporal trend among all drivers (Fig. 1B). Snowmelt exhibited no long-term trend, but an expected seasonality. Precipitation patterns were stochastic, with no detectable trend. Modelled topsoil moisture (first 30cm layer) showed a strong seasonality consistent with summer droughts, with additional multi-year variability and pronounced minima during the droughts of 2003 and 2019-2022 (29, 30). It also showed a linear decrease through time (Fig. 1B).

**Fig. 1.**
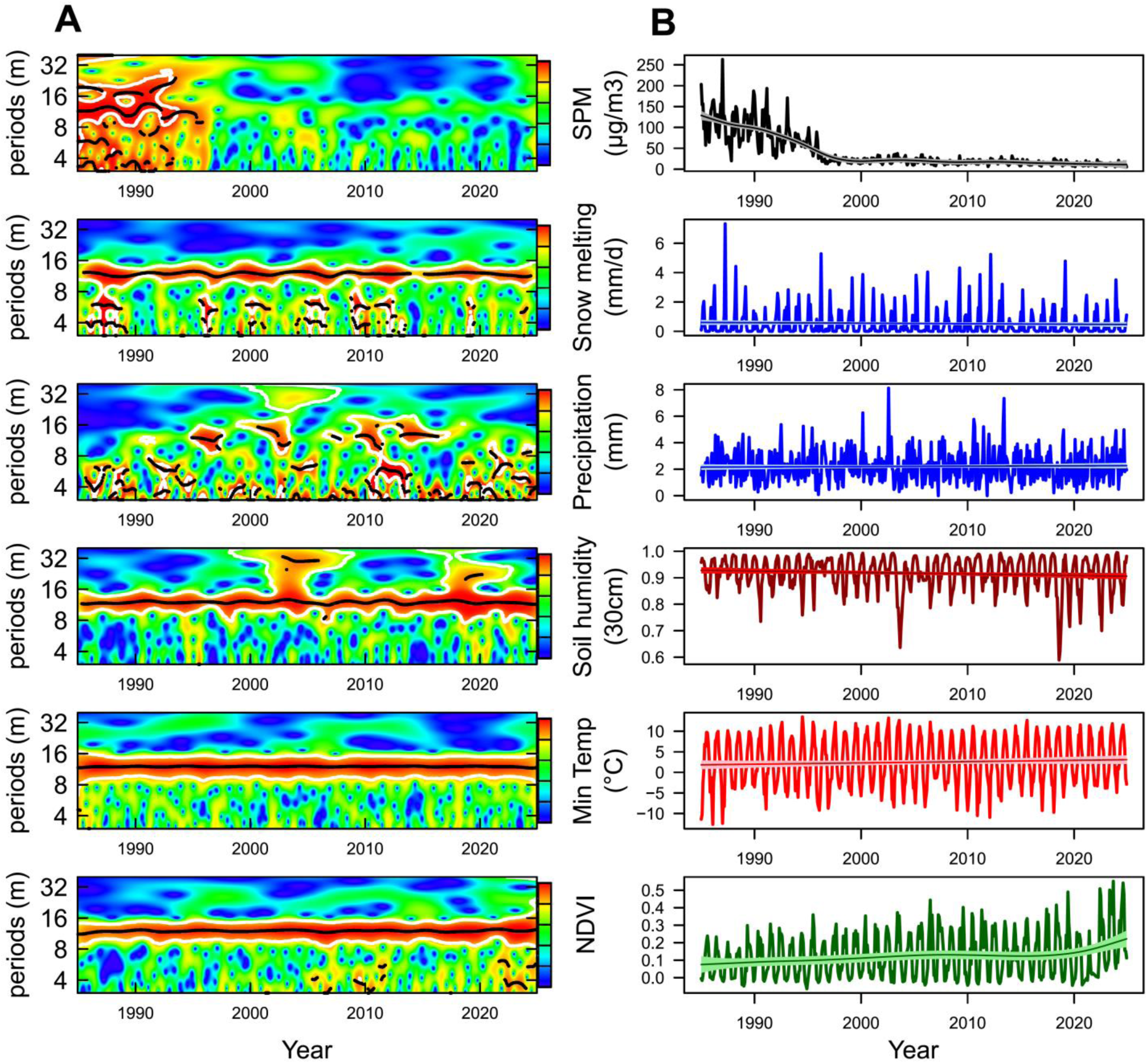
Four decades of atmospheric pollution, meteorological, and hydrological changes. (A) Wavelet power spectrum analysis showing the recurrence patterns in time series for each driver of global change considered. Gradient intensity from blue to red indicates the spectral power, highlighting periods with the greatest temporal variability through time (x-axis), while the period of recurrence frequency of the cycles are indicated vertically(y-axis, in months). Areas surrounded by white are statistically significant areas of the spectrum. (B) Corresponding time series in their natural units, with fitted smoothed GAMs, the main fit, and 95% credible intervals. *SPM GAM*; e.d.f. = 9.8; F = 121.1; p <0.0001; Deviance explained = 76.1%; *Snowmelt GAM*; e.d.f. = 1.0; F = 0.837; p = 0.361; Deviance explained = 0.18%; *Precipitation GAM*; e.d.f. = 1.0; F = 0.353; p = 0.553; Deviance explained = 0.07%; *Soil humidity GAM*; e.d.f. = 1.0; F = 4.673; p = 0.0311; Deviance explained = 0.97%; *Minimum temperature GAM*; e.d.f. = 1.0; F = 1.795; p = 0.181; Deviance explained = 0.38%; *NDVI GAM*; e.d.f. = 4.641; F = 4.847; p < 0.001; Deviance explained = 6.55%.

Normalized Difference Vegetation Index (NDVI) ranged from zero to negative in winter (dormant vegetation) to positive in summer, reflecting seasonal vegetation growth, or “greenness”. NDVI steadily increased from 1985 to 2015 (Fig. 1B), corresponding to the early-forest succession of sparsely planted Colorado spruce trees (SI Fig. S2). The increase in NDVI in the last decade may reflect the replacement of non-native Colorado spruce with native Norway spruce which may have had higher growth rates from 2015 to 2025 (Fig. 1B).

### Long-term community dynamics and critical transition

To model population dynamics, we used both a multivariate (dynamic) Bayesian generalized additive model (M(D)GAM) (48) and wavelet power spectrum (WPS) analysis (49). During high pollution loads in the 1980s–1990s (Fig. 1B), the largest herbivore *Microtus agrestis* (∼20–40g adults) was dominating abundances, with quasi-cycles of large amplitudes at multiple recurrence frequencies of ∼1.5, ∼2.5, and ∼4.5 years (Fig. 2A; SI Fig. S3), whereas *A. flavicollis*, the largest omnivore in the community (∼20–40g adults), showed cycles of relatively low amplitude (Fig. 2A; SI Fig. S3).

**Fig. 2.**
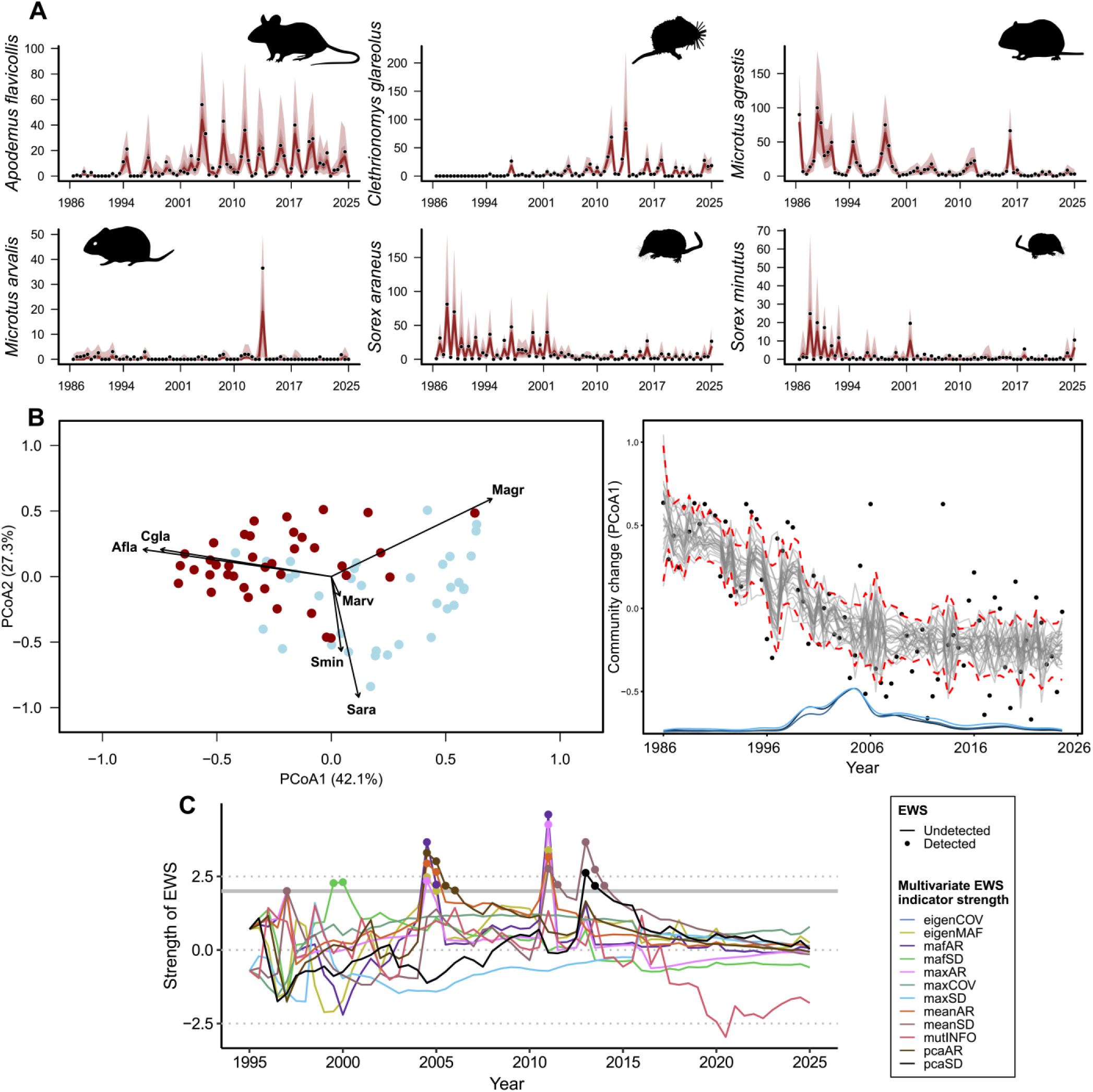
Small-mammal community dynamics and critical transition. (**A**) Species exhibited cyclic population dynamics. Observations are as black dots, and predictions with point estimates and 95% credible intervals are shown in dark red. Silhouettes of each species are from ‘philopic.org’ (public domain, except for *Microtus arvalis;* credit to “T. Montessuit”). (**B**) Ordination of the community on a Principal Coordinate Analysis (PCoA) biplot, with in blue, time steps before 2005, and dark red, time steps after 2005 (left panel). Modelled trajectories (grey lines and 95% credible intervals in dashed red lines) of a multiple change point model including temporal autocorrelation, with in blue, the posterior distributions of the identified critical transition (right panel). (**C**) Multivariate Early Warning Signals (EWS) analysis. EWS are depicted as colored dots over years of the time series, with colors relative to several EWS indicators.

Since the mid-2000s, *A. flavicollis* was the species dominating abundances in the assemblage, exhibiting large amplitude population quasi-cycles with multiple recurrence at ∼1.5 and ∼3 years frequencies, thus potentially showing period doubling. *C. glareolus*, another omnivore (∼20–25g adults), showed quasi-cycles at ∼2 years frequency, with the largest amplitudes in the 2010s (Fig. 2A; SI Fig. S3). Both *Sorex spp.* insectivore shrews exhibited seasonal quasi-cycles of ∼1 year frequency, with potential signs of period doubling at ∼2 and ∼4 year frequencies, with the largest amplitudes observed on the first half of the time series (Fig. 2A; SI Fig. S3).

To interrogate the eventuality of a critical transition, we first reduced the dimensionality in community composition (Fig. 2B). Our Principal Coordinate Analysis (PCoA) confirmed that the community composition for the first half of the time series was dominated by *M. agrestis* and *Sorex* spp. abundances, with omnivores dominating abundances in the second half. With a multiple change-point Bayesian model, we detected a critical transition in 2005 (1998–2024; high probability density ‘HPD’ intervals at 95%; Fig. 2B). Early warning signals (EWS) of a critical transition occurred in 1997, followed by marked multi-indicator signals in 2000 and 2005 (Fig. 2C). Therefore, marked signs of a community transition were observed in 2005, seven years after the inflection of atmospheric pollution (Fig. 1B).

### Delayed density-dependence

We used the M(D)GAM to examine the predicted impacts of sudden increases the abundance of one species on the system. Simulated shock impulses in *A. flavicollis* abundance generated strong self-regulation as well as effects on all other species: *C. glareolus*, *Microtus spp.,* and the small-bodied (∼2–10g adults) *Sorex spp.* (SI Fig. S4). Density-dependent responses of all species to shock impulses of *A. flavicollis* abundances were delayed and “wiggly” up to four years after pulse perturbations. Applying this approach to all species enabled variance decomposition of impulses, suggesting that *A. flavicollis* was relatively unaffected by shocks from other species, but had a large effects on other members of the community (SI Fig. S5).

### Links between global change drivers and population dynamics

Consistent with our hypothesis, omnivores and in a lesser extent, insectivores, had delayed declines to increases in SPM (SI Fig. S6). These negative effects were strongest at 1- and 3-month lags, and matched with lagged positive responses in herbivores at 2- and 4-month lags (SI Fig. S6), perhaps as a result of competitive release (apparent facilitation). Using convergent cross mapping (CCM) from EDM (44), we confirm that SPM was causally-driving the two omnivores, and one of the main drivers controlling *M. agrestis* and *S. minutus* (SI Fig. S7). Vegetation index (NDVI) and minimum temperature had strong causal effects on shrews (SI Fig. S7).

### Atmospheric pollution induced chaos and strengthened interactions

Lyapunov exponents (LE, λ) mathematically relate to the presence, absence, and intensity of instability, with null or negative values indicating stability (the system would converge to equilibrium after a perturbation), and with positive values indicating instability (the system would diverge exponentially after a perturbation) (1). The community fluctuated at the edge of chaos (Fig. 3A), with rare negative LEs and common periods of instability (*MCMC* posterior mean λ: 0.80, 95% credible intervals: 0.57–1.01; Fig. 2A). We detected a causal effect of SPM on chaos (*ρ* = 0.44; *p* = 0.005; Fig. 3B), which we did not find for other drivers (SI Fig. S8).

**Fig. 3.**
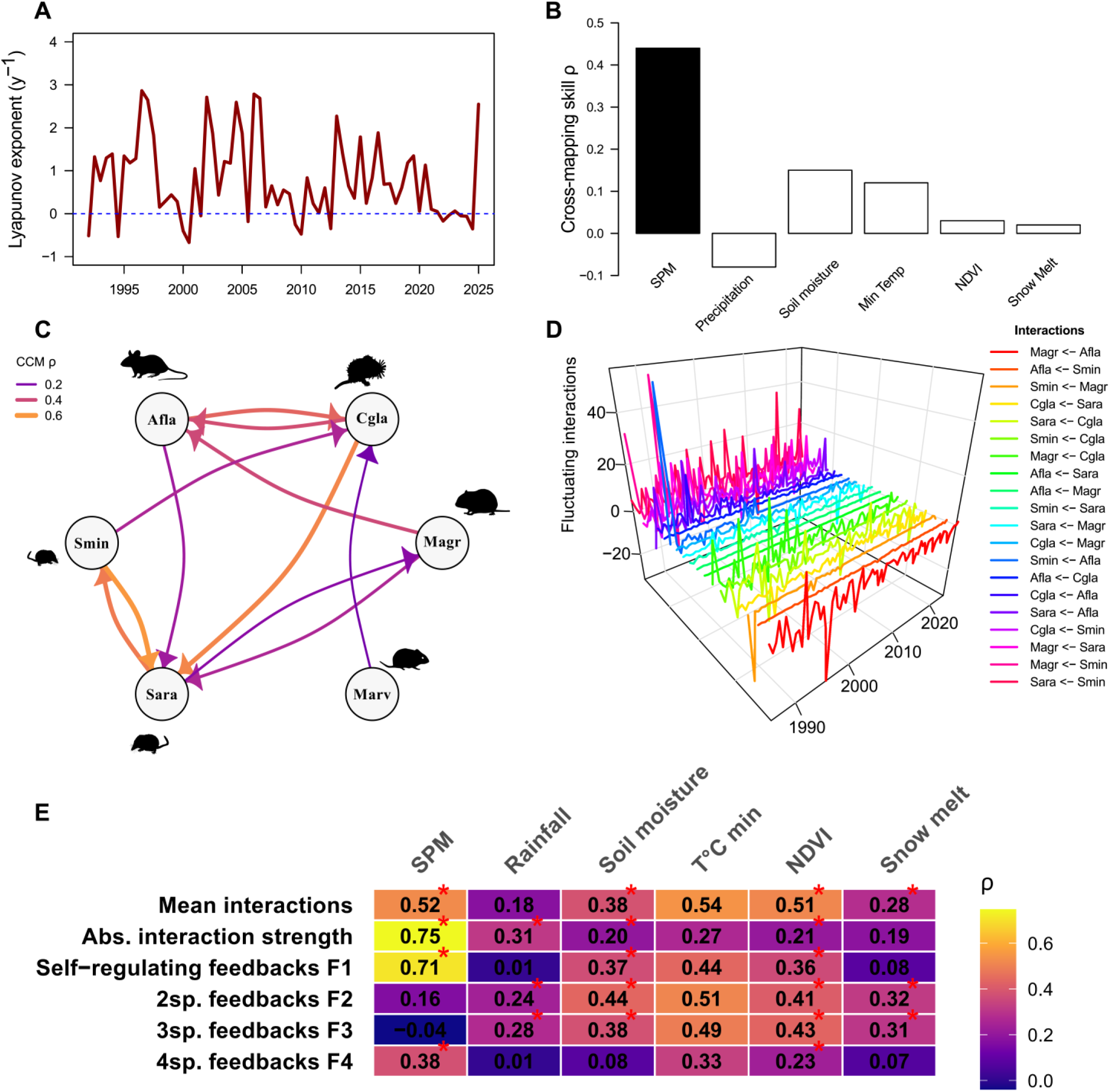
Chaos and interaction strengths were driven by atmospheric pollution. (**A**) Lyapunov exponents through time. (**B**) Causal effects (forecast skill *ρ*, the Pearson’s correlation coefficient between predicted and observed time series at maximum library size) of environmental drivers on Lyapunov exponents; the black bar shows the significant effect of SPM. White bars show non-significant effects relative to surrogate distributions of seasonal null time series. (**C**) Parsimonious network (from CCM) showing the strength and direction of causal links in the network, with the color and size of arrows proportional to the forecast skill *ρ* (“Afla”: *A. flavicollis;* “Cgla”: *C. glareolus;* “Magr”: *M. agrestis;* “Marv”: *M. arvalis;* “Sara”: *S. araneus, and* “Smin”: *S. minutus*). (**D**) Time-varying species interactions for the full network obtained from S-maps coefficients. (**E**) Causal effects tested by CCM (predictive skills ρ), with heatmap of the strength of causal effects for each environmental driver on network interaction and feedback loop metrics. Red asterisks indicate significance (observed *ρ* > 95% of 1000 null seasonal surrogates, i.e., *p*<0.05).

We used CCM (2) to reconstructed a parsimonious network of interactions (Fig. 2C). In this network, we identified twelve feedback loops of interactions, ranging from one-species (self-regulation) to four-species loops, and explored their stabilizing/destabilizing properties empirically (50) (SI Appendix 1). We used multivariate S-maps among 20 species pairs to model time-varying interactions and feedbacks (Fig. 3D; SI Appendix 1; Fig. S9a-b; and Fig.S10). During the polluted period, the network exhibited abrupt changes in interaction signs and strengths (Fig. 3D and SI Fig. S9a), marked by a few strong negative feedbacks of the four-species loops (F4) immediately mirrored by destabilizing positive feedbacks of the two- (F2) and three-species loops F3 (Fig. S9a). Around the identified critical transition (∼2005), feedback loops often showed an even number of negative interactions leading to positive self-amplifying feedbacks (SI Fig. S9a,b). The self-regulating feedbacks indicated by negative F1 values spiked just before 2005 (SI Fig. S9a).

Atmospheric pollution was one of the main drivers of interactions’ signs and strengths, and affected strongly the feedbacks of one- (F1) and moderately the feedbacks of four-species (F4) loops (Fig. 3E). Soil moisture and NDVI had moderate effects on interaction metrics (Fig. 3E). Overall, this confirms that pollution, in addition to droughts and changes in vegetation, played an important role in dynamically-reshaping interactions (Fig. 3E; SI Fig. S9).

Finally, we explored *how* environmental drivers affected the interaction metrics (SI Fig. S11), and whether and how these changes in interactions related to chaos (SI Fig. S12 and S13). SPM and NDVI positively influenced self-regulation feedbacks (F1) (SI Fig. S13). Feedbacks F2 and F3 were positively influenced by soil moisture and NDVI (SI Fig. S13), perhaps reflecting ecosystem productivity. On the contrary, the feedbacks F4 were negatively related to SPM levels and soil moisture (SI Fig. S11). NDVI and SPM increased the mean signed interactions and the mean absolute interactions in the network (SI Fig. S11). Finally, Lyapunov exponents (LE; λ) were positively influenced by the F2 and F3 feedbacks, and by the mean absolute interaction with SPM levels (SI Fig. S12-13).

### Forecasting with or without key environmental drivers

Adding atmospheric pollution in multivariate S-maps led to improved forecasting from 1.5 up to 2.5 years, and forecast skills from 0.1 to 0.6 *ρ*, especially for the pollution-sensitive *A. flavicollis* (SI Fig. S14). Minimum temperature, when added to S-maps, was detrimental for forecasting *S. araneus* dynamics, as expected. Adding SPM in addition to NDVI showed little forecasting improvements (SI Fig. S15).

## Discussion

Here, we show how concomitant environmental changes contributed to non-equilibrium dynamics and fluctuating interactions in a community of small mammals. Our analyses suggest that atmospheric pollution negatively affected the omnivorous species, strengthened species interactions in the network, contributing to chaos. The decline in pollution levels induced a qualitative change in the system’s dynamics, with several indicators of a critical transition detected in 2005, only several years after the rapid pollution decline. This result suggests that the recovery of communities following perturbations can be abrupt, rather than on the form of a smooth recovery. The resulting omnivore-dominated regime was structured by weaker interactions. Therefore, our four-decade study in the air-polluted Black Triangle offers unique insights into the mechanisms leading into community instability (38) in the Anthropocene.

Dynamical instability, or chaos, is expected in a wide parameter space in multi-species theoretical models (51), even without environmental stochasticity nor periodic forcing. In our small mammal’s network, field vole and yellow-necked mouse abundances oscillated at low frequency but large amplitude, while shrews fluctuated at high seasonal frequency. The resonance between interacting species with mismatching abundance frequencies can theoretically cause chaos (52). Shrews, which are amongst the smallest body-sized mammals in Europe, had their population sizes strongly forced by seasonal vegetation growth (NDVI) and minimum temperatures (CCM with *ρ* up to 0.95). Shrews’ fast metabolic rates constrain them to forage continuously on soil-dwelling invertebrates, and as shrews do not hibernate, they are strongly forced by seasonality and arthropod productivity which were probably captured by minimum temperature, soil moisture, and NDVI. Our loop analysis showed that the F2 and F3 feedbacks were destabilizing and were mostly driven by those same environmental drivers. F2 and F3 feedbacks were linked to chaos, suggesting that seasonal changes in vegetation (e.g., early-forest succession) and soil moisture were contributing to a first route to chaos.

Lead (Pb) associated to atmospheric deposition likely contributed to nearly suppress the two pollution-sensitive omnivores (with effects of SPM delayed from between several months to two years; SI Fig. S6 and Fig. S20). The omnivores’ abundances, however, exhibited a few small peaks (Fig. 2A) during the polluted-era, suggesting they maintained baseline abundances in the landscape. The yellow-necked mouse is an aggressive competitor [e.g., a “killer mice” (53, 54)], making it probably keystone to the network (45, 46) and to the pollution-induced instability that we observed. Moreover, Pb is known to increase aggressivity in rodents (55), and could have amplified interference competition from the few remaining omnivores, strengthening their per capita effects despite their low densities. This could have resulted in a strengthening of interactions, the latter causing chaos with a delay of 1.5 years (Fig. S12). Following peaks of pollution, the shrews and the field vole — freed from the control by omnivores and fueled by the abundance of grassland patches — entered overcompensatory feedbacks, overshooting their carrying capacity in the form of “boom- and-bust” dynamics (38). In addition, massive nitrogen deposition during the pollution era (up to 25 Kg ha^−1^ yr^−1^) (56), by fertilizing the grass could have boosted the large cycles of the herbivore *M. agrestis*, adding a new perspective to the discussion of their dampening oscillations by Cornulier et al. (38, 41).

The F1 and F4 feedbacks were often negative, but not sufficiently negative enough to counterbalance or “brake” on the strongly destabilizing feedbacks F2 and F3 (SI Fig. S9a) (50). A flickering between negative and positive feedbacks, strong and nonlinear feedbacks, and overall positive feedbacks, theoretically lead to chaos and bifurcations (14, 57). For instance, after 2005, when we identified the onset of a critical transition (Fig. 2B), the sum of all *F* feedbacks at any level of *k* species (i.e., F1, F2, F3, and F4) were net-positive (SI Fig. S9a) (50, 14), thus featuring the destabilizing positive feedbacks underlying instability and critical transitions (47). Moreover, we found that atmospheric pollution reduced self-regulating effects of the species abundances on themselves, promoting unstable dynamics. During the low-pollution era, the omnivores recovered, and relatively to the polluted period, dampened network instability (45, 46). Our insights therefore add to other ecological examples (2, 18) of environmental changes causing interactions to change in magnitude and sign (competition to facilitation).

Therefore, our results are corroborating ecological theory. First, with a marine trophic network model, Cunico et al. (58) showed that omnivory is stabilizing (limiting chaos), confirming earlier theory (45). Second, they found that increased interaction strengths are destabilizing (58), a result that is common to our empirical study and the one of Ushio et al. (2), and several theoretical works (59, 58). Third, we showed that fluctuating chaos was cross-mapping with the positive feedbacks by two- and three-species loops (i.e., F2 and F3 “caused” chaos). This is theoretically sound, as in networks of mutualism and/or competitive interactions (as here), two- and three-species loops are predicted to be the main destabilizing factors (59). Positive feedbacks are also necessary for ecosystems to exhibit multiple stable basins, and/or transients between these multiple stable states (21). That said, we are conscious that statistical modeling of empirical data cannot fully confirm the deterministic “truth” of a system’s behavior. This is the case since real-world data include both stochastic noise and observation error, natural complexity, and unobserved factors. Therefore, modeling (indeed, both theoretical or statistical) stays a mere approximation of reality (1). For instance, loop analysis has been designed for systems at steady state, and is thus often reserved for the analysis of theoretical models (50). However, as expressed in Paige et al. (21), we think that empirical statistical analysis of feedback loops is informative on the processes leading ecosystems to be unstable and exhibit critical transitions.

## Conclusion

Small mammal dynamics have fascinated several generations of ecologists and have been pivotal to our knowledge of cycling populations (60, 38), chaos (11, 37), transients (61) and community changes (35, 36, 39). So far, the causes of these phenomena have been attributed to predator-prey cycles (37), masting cycles (36), climate (62), and land-use [e.g., afforestation, urbanization (39, 62)]. *In addition to these and potential unobserved factors*, our results show that an overlooked factor —atmospheric pollution— was detrimental to the “clock mechanics” of a small mammal community. By synthesizing four decades of changes, we show that identifying the environmental drivers of ecological instability is essential for attributing causes of ecological changes (63), understanding the underlying mechanisms of these changes, and improving ecological forecasts of chaotic dynamics (34, 5). Atmospheric pollution conveying toxic elements is likely to decline even further in Europe (Zero Pollution Action Plan to reduce air pollution to non-harmful quantities for human and non-human life by 2050), offering opportunities to assess the effectiveness of policies. Many regions, however, continue to suffer from severe atmospheric pollution [99% of human populations are experiencing bad air quality, the leading environmental cause of mortality in the world, WHO (64)], and we need to assess urgently the effects of emergent contaminants (65) amid climate change. We hope that our study will encourage multidisciplinarity, a better acknowledgement of pollution as a global change driver (24, 33, 25), and the study of long term dynamics of entire ecosystems (26).

## Methods

### Study site

Our study spans 39 years (1986–2024) in the Ore Mountains (820 - 860 m a.s.l.), in NW Czech Republic (50.7°N, 13.6°E). From the 1960s to the 1990s, notoriously-high emission loads from brown coal and chemical plants caused acidic rains and metal(loid) deposition, for instance, by lead (Pb) (SI Fig. S1), resulting in diebacks of Norway spruce (31). After clear-cuts of the dead stands in the late-1980s, European birch, European mountain ash, larch, and the non-native Colorado blue spruce were sparsely planted (32). From the 1980s until 2010, the landscape developed into a mosaic of young stands and open areas, with a ground layer dominated by hairy reed grass over the study period. During the 2010s, the widespread blue spruce plantations were replaced by native Norway spruce after a fungus (*Gemmamyces piceae*) damaged the blue spruces (SI Fig. S2).

### Small mammals sampling

We conducted semiannual (beginning of June and October) snap-trapping from 1986 to 2024, a reliable method for estimating the abundances of ground-dwelling small mammals (37, 66, 36). We conducted trapping in three 1-ha plots spaced by ∼2km, each plot divided in an 11×11 grid with individual snap-traps spaced 10-m apart (121 traps per plot). In each trapping session, we operated traps for three days, checking them and resetting them each morning. Roasted flour mixed with bacon was used as bait. To avoid any observational bias, autumn plots were adjacent to spring plots (SI Fig. S2).

### Monthly environmental data

For our measure of thermal conditions, we used monthly mean minimum temperature as it constrains thermoregulation of small homeotherms (67). Precipitation and snow cover are also critical to rodents, affecting hibernation, habitat and resource use. We focused on snowmelt, as we hypothesized as well as describing the amount of snow in winter and subsequent water release in spring, that snowmelt could release the deposits of toxins accumulated in the snow during winter. NDVI described early-forest succession and vegetation growth (68). Soil moisture in the top 30cm of soil is linked to invertebrate activity, ecosystem productivity, and major drought events (30, 29). We added to these monthly data suspended particulate matter (SPM), comprising a wide size range of dust particles (<100µm), probably associated with toxic elements, such as lead (Pb) (SI Fig. S1). Full details on monthly environmental data acquisition and modeling are given in SI, Appendix 2.

### Data analysis

#### Data preparation

Over the entire period, we trapped a total of 3466 individuals belonging to 15 species. For our analysis, we used data on six abundant species (98.24% of total abundances). We used R (version 4.5.2.) (69). Within each trapping period, the numbers of individuals trapped across the three plots (SI Fig. S16) were summed by species. Our time series analysis required equal temporal spacing of data, and so we computed these equidistant data (6-months interval) from June and October observations using cubic Hermite interpolation (to 15^th^ June and December of each year) (6) (SI Fig. S17).

#### Time series analysis of environmental drivers

We described trends for each driver using wavelet power spectrum (WPS) (70). WPS consists of continuous Morlet transforms to identify patterns in the dominant frequencies (i.e., the power) in time series [R package *‘WaveletComp’* (49)]. We modelled long-term trends in each external driver with generalized additive models (GAMs) [R package ‘*mgcv* ‘(71)], with a smooth of time (*k*=50 basis functions). By using REML optimization, the GAMs naturally penalized high-frequency seasonal fluctuations down to low effective degrees of freedom (e.d.f = 1–9.8 in all cases), modeling inter- annual trajectories.

#### State-space modeling of community dynamics

We first jointly modeled the dynamics of species in the small mammal community using a Bayesian multivariate (dynamic) state-space generalized additive model (MDGAM) [‘*mvgam*’ R package (48)]. This model separates the underlying latent ecological process from observation noise. This accounts for sampling error and imperfect detection on the observation scale, while capturing the shared latent trend and group-specific (here, trophic guilds) trends. This multivariate structure allowed us to assess density-dependence, providing complementary insights into community assembly. We also tested for delayed effects of atmospheric pollution at the guild level (see SI Appendix 3).

#### Critical transition and Early Warning Signals (EWS)

We tested for a single critical transition in community dynamics that we hypothesized to exist following the decline in air pollution. We first used a Principal Coordinates Analysis (PCoA) on Hellinger-transformed abundance data (to avoid the double zero issue), to reduce the dimensionality of the six species community to two dimensions. We retained the time series of PCoA axis 1 for critical-transition analysis. To account for temporal autocorrelation in our community composition time series (PCoA1), we modeled structural transitions in the PCoA1 scores using Bayesian multiple change-point analysis [“*mcp*” in R (72), linked to JAGS]. The first segment was defined as a linear trend with a second-order autoregressive error structure “AR(2)”. The second segment captured a relative change in slope combined with a first-order autoregressive structure “AR(1)”. Parameters were estimated using 30,000 iterations across three chains, and default priors. The transition timing was reported using the posterior mode and its 95% highest posterior density intervals (HPD). For all parameters estimated using Bayesian models, Monte Carlo Markov Chains (MCMC) mixed well, as indicated by Rhat values close to one.

We assessed early warning signals (EWS) (i.e., changes in variance-covariance, autocorrelation, and/or critical slowing down) of a critical transition using multi-index EWS [R package ‘*EWSmethods*’ (73)]. We considered a significant EWS as two consecutive warning signals above the 2σ threshold. Confidence for an EWS increased with the number of indicators exhibiting warnings (74).

#### Empirical Dynamic Modeling (EDM)

EDM (44) is a nonparametric “equation-free” approach based on Taken’s theorem, reconstructing the state-space of a nonlinear and potentially chaotic system from one or more time series of that system. This allows testing for causality (44), forecasting, and recovering time-varying interactions (18).

##### a. Time-varying chaos

We described time-varying chaos using local Lyapunov exponents (LLEs) (15, 6, 2). We define our measure as unconditional LLEs or “apparent” chaos, where both stochastic and deterministic processes contribute to realized stability or chaos [*sensus* Munch et al. (1)]. We used a Jacobian- based estimation of LLEs (1) on our six-species time series from multivariate S-maps (2) and a time window of 15 time steps [i.e. 7.5-y; R package ‘*EWSmethods*’ (73)]. We chose the Jacobian method as it has been shown as the most robust against noise (8). By using unconditional LLEs, we avoid making assumptions about exogenous drivers in the estimation of chaos itself (unlike conditional LLEs), and are able to test independently the potential causal drivers a posteriori, for instance, using convergent cross mapping (CCM) (44).

##### b. Testing causality between time series: convergent cross mapping (CCM)

CCM evaluates whether historical values of a variable *X* can reconstruct the dynamics of another variable *Y*. This is based on an extension of Taken’s theorem, which posits that if two variables are causally linked, their univariate attractors are invariant to the true deterministic attractor, and are thus topologically-invariant from each other (44, 18). In that case, the two attractors of *X* and *Y* map “one-to-one” in the state space of the attractor/manifold. Since *X* is causally influencing *Y*, the dynamics of *X* are imprinted in *Y*, and thus, we can predict *X* with *Y* in choosing the best embedding dimension *E*. We say that *Y* crosses map *X*, or *X* causes *Y*. The direction of the cross mapping is opposite to causality: if a predator’s dynamic “causes” the one of a prey, the dynamics of the predator get “printed” into prey’s ones, so we can predict predator’s dynamics from prey’s ones (18).

Before CCM, we identified the optimal embedding dimension *E* using simplex projection, which reconstructs the attractor/manifold using lags from the response variable. For instance, an attractor from *Y* with *n* embedding dimensions (*E* = *n*) is represented by *n*+1 lagged vectors: *Y_t_* =<*Y_t_*, *Y_t-1_*, *Y_t-2_* (…) *Y_t-n_*> (18, 44). For each variable, we identified *E* with the highest prediction skill (*ρ,* the Pearson’s correlations between the observed and predicted values for that *E*).

We quantified convergence (a condition to distinguish causation from correlation) (18, 44) when *ρ* increases as a function of total library size *L* (the length of the time series minus *E*). Convergence occurs as the data to reconstruct the manifold increase, predictions from closest neighbors in the state space improve (18, 44). Once we identified the best E, we re-ran each CCM with delays *Tp* from –3 to 3, and kept the *Tp* value optimizing *ρ_obs_* at *L_max_*. We considered causation to have been identified when 1) the observed skill *ρ_obs_* at *L_max_* was > 0; 2) *ρ_obs_* at *L_max_* > *ρ_obs_* at *L_min_* (convergence); and 3) *ρ_obs_* at *L_max_* > 95% of the *ρ_null_* from a null distribution of a thousand seasonal surrogates (2, 18). We applied CCMs to (*i*) infer causal effects of environmental drivers on biological dynamics (e.g., populations, chaos), and (*ii*) infer causal interactions among species to infer a parsimonious network of interactions. From this network, we identified twelve feedback loops of levels one- (F1), two- (F2), three- (F3), and four-species (F4) interactions (50) (SI Appendix 1). Convergence plots, *E*, delays *Tp*, and *p*-values for all CCMs are available as supplementary information (SI Fig. S8, S12, S18, S20, S21).

##### c. Time-varying interactions with S-maps

We reconstructed time-varying species interactions using multivariate S-maps (sequential locally weighted global linear maps) (75, 76, 2, 18). S-maps estimate local linear approximations of the reconstructed state-space or manifold, recovering information about how species interactions vary through time (magnitude and direction). Multivariate S-maps apply the same rationale as above, but using multiple variables to reconstruct a multivariate manifold. S-maps weigh library vectors exponentially by their distance to the prediction vector in the multivariate manifold before using linear regressions to make a forecast; library vectors closest to the prediction vector have the greatest weight. To predict a point *x\** in the manifold, each library vector or point *x_k_* in the attractor is given a weight:

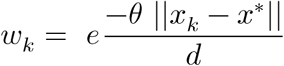

Where ||*x_k_* – *x*\*|| is the Euclidean distance between the target and library vector, and *d* is the average distance among library vectors in the manifold (18, 75, 76).

For all species pairs (20 interactions), we extracted time-varying coefficients from the S-map’s Jacobian matrix (18, 44, 76). We included the interaction of *M. arvalis* on *C. glareolus* in the S-map of the latter to stabilize other species interactions on *C. glareolus*, but as *M. arvalis* exhibited a strong peak of abundance that was poorly explained by environmental drivers, we disregarded this vector from subsequent analyses (*M. arvalis* on *C. glareolus*). Each multivariate S-map was optimized to its best nonlinearity parameter *θ* that maximizes the predictive skill but setted *θ_max_* = 15 to avoid hyperlocality (see SI Fig. S19). S-maps approach a global linear model when *θ* equals zero, and a nonlinear state space model when *θ* > 0 (nonlinearity). We computed metrics reflecting interactions in the network from the Jacobian derivatives of the interspecific interactions: the mean of signed interactions and the mean absolute interaction strength (2). We also computed the feedback values for the 2-species, 3-species and 4-species loops (50) (see SI Appendix 1). We modelled self- regulating feedbacks (F1) from one-species feedback loops using separate univariate S-maps. To ensure the self-regulating intraspecific coefficients are not overestimated and stay comparable with interspecific ones, we removed – 1 to each of these one-species time series (77). We used these metrics to describe whether environmental drivers caused changes in interactions in the network with CCMs, and for qualitative interpretations, rather than for a rigorous loop analysis using Levin’s stability criteria. Our rationale was that our system exhibits non-equilibrium dynamics (and a rigorous stability analysis requires a steady state), and our self-regulating coefficients were not computed together with the interspecific coefficients. However, we believe that a loop analysis on empirical data still provides valuable information on the mechanisms resulting in instability and abrupt changes (21).

##### d. Exploring the mechanisms of changes in interactions and chaos

To explore *how* environmental drivers influenced community instability, and in turn, time-varying Lyapunov exponents, we used multivariate S-maps graphically displayed with surface plots.

Response surfaces consisted of 100 bootstraps of S-map predictions on a 100×100 grid of two drivers, with an exclusion radius of 12 to smooth the interpolations [following Merz et al. (76)].

##### e. Forecasting with or without key predictors

We performed leave-one-block-out forecasting (34) using multivariate S-maps, with or without key environmental drivers. S-maps were set-up with an exclusion radius of 2 (i.e., excluding two time steps before, and two time steps after the time step to be predicted), at increasing time horizons (from *Tp* 1 to 5, i.e., from 6 months to 2.5 years). Forecast accuracy was quantified using Pearson’s *ρ* between observed and predicted values. We compared models including environmental drivers to baseline models, assessing the contribution of drivers to forecast ability through a better unfolding of the state space manifold (34). All S-maps specifications and fitting parameters are available (SI Table S1). For CCMs and S-maps, we used the R package ‘*rEDM*’ (44).

## Declaration on AI use

We used gen-AI (free versions of OpenAI ChatGPT [GPT-4o mini/GPT-5/6 Luna] and Google Gemini [Gemini 3 Flash/Flash-Lite] between Jan-26 and Oct-26) to assist us designing some R functions for data handling, repetitive calculations (CCMs and S-maps), and writing plotting code. All hypotheses, conceptualization, workflow of data analysis, data analysis rationale, interpretation of the results, their validation, and writing, were performed by the leading authors based on available resources and literature, and thus, the authors are accountable for the work reported here.

## Data availability

The biological and environmental datasets used for this study, along with the R code to reproduce the statistical analyses and graphics are available at Figshare [https://doi.org/10.6084/m9.figshare.34906974] (78). A web view of the R code and outputs is available.

## Ethical Statement

This research was conducted in accordance with national and international ethical standards and Czech national legislation. Small mammal trapping was conducted as part of authorized wildlife monitoring activities in the study region under a permit issued by the Ministry of the Environment of the Czech Republic (No. 71735/ENV/16-3580/630/16).

## Conflict of interests

The authors declare no conflict of interest.

## Authors’ contributions

**T.R.**: Conceptualization, research questions, statistical analyses, fieldwork, writing the first draft and editing until its final form. **V.B., K.S., F.S., J.Z.**: Project initiation, coordination, long-term fieldwork, data acquisition. **O.R.**: Modeling, acquisition, and processing of meteorological, hydrological, and vegetation-index data. **J.S.**: Fieldwork, data acquisition. **R.S.**: Fieldwork, writing, data acquisition. **A.S., V.C.**: Laboratory analyses of lead concentrations. **W.M.H.**: Critical inputs on an early draft, editing. **M.Z.**: Conceptualization, secured funding, project coordination, data acquisition, curation and validation, fieldwork, supervision. All authors agreed on the submitted version. We dedicate this manuscript to the zoologist Jan Zima, one of the initiators of this long-term monitoring.

## Acknowledgements

We are thankful to all students and volunteers involved in fieldwork over the years: without your support, this work could not exist. The analyses and preparation of this manuscript were supported by the Czech Science Foundation (GAČR), under the Grant No. 25-17362S: “*Small mammals as key indicators of sublethal impacts of air pollution and metal(loid) deposition in terrestrial ecosystems (POLLCHAIN)*”, awarded to Markéta Zárybnická (PI).

## Notes

https://doi.org/10.6084/m9.figshare.34906974

### Competing Interest Statement

The authors have declared no competing interest.

### Summary of Updates

Minor text improvements and add the analysis of environmental changes in the main text

