## Supplementary Information for "Atmospheric pollution contributed to destabilizing a small mammal community amid four decades of environmental changes"

### Appendix 1: Loop analysis

Feedback loops have been the object of early attention in ecology (1). Richard Levins in 1974 (1) posits the basics of loop analysis, in that the feedback of *m* disjoint loops in a given combination of loops of *k* species would equal to $F_k$ as follows:

$$F_k = \sum_{m=1}^{k} -1^{m+1} \sum L(m,k)$$

With *L(m,k)* the product of the loop coefficient values for a combination of *m* disjoint loops that contain *k* species. Two loops are disjoint when they do not share any species.

When $F_k$ is positive, the loops $L_k$ exert positive feedbacks (amplifying perturbations, promoting instability), and when $F_k$ is negative, the loops $L_k$ exert negative feedbacks (reducing perturbations, promoting stability). Therefore, exploring the dynamical behavior of feedback loops can be insightful to our understanding of instability and critical transitions in our small mammal assemblage.

In our parsimonious network (obtained from CCMs), we identified twelve loops at $k = 1$ (five loops), $k = 2$ (three loops), $k = 3$ (two loops), and $k = 4$ (two loops) (Figure A1).

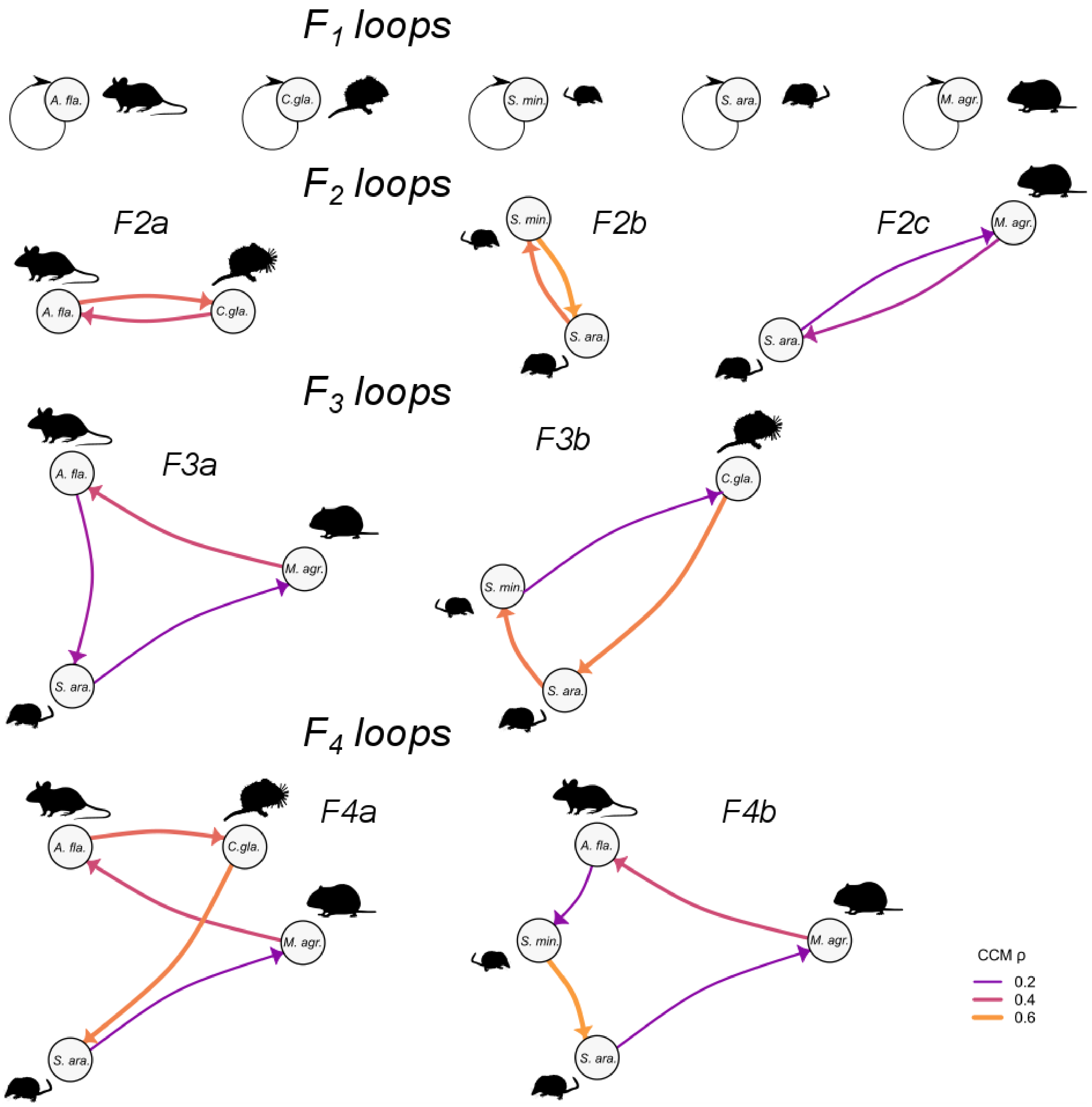


**Fig. A1**: The twelve feedback loops in our system.

In these conditions, F1 is equal to the sum of self-regulating feedbacks for species *ij* at each time step.

F2 is equal to the sum of the products of the two-species loops (LF2a + LF2b + LF2c).

F3 is equal to the sum of the products of the three-species loops (LF3a + LF3b).

F4 is obtained from the four-species loops and from two pairs of disjoint two-species loops (F2a / F2b and F2a / F2c; Figure A1), with for these disjoint loop pairs, $m$ = 2 so that $-1^{m+1}$ in the general expression gives $-1^{2+1} = -1$, therefore we have F4 = (LF4a + LF4b) – (LF2a × LF2b) – (LF2a × LF2c).

Theoretically, and at steady state, the system can be stable only if all feedbacks at $k$ levels are negative.

In addition to visualize temporal changes in F1, F2, F3 and F4 values, and test if any environmental driver could cause variation in feedbacks at these levels (e.g., with CCMs), it can also be informative to qualitatively describe changes in the conditions for instability (positive feedbacks) and stability (negative feedbacks).

For a two-species loop, if both interactions are negative or positive, their product is positive. It requires a positive and a negative interaction (exploitation) for the loop to be stabilizing (negative product). For a three-species loop, the feedback is positive when three interactions are positive or when two interactions are negative and one is positive. The feedback is negative if three interactions are negative, or if one interaction is negative and two are positive. For a four-species loop, the feedback of the loop is positive when all interactions are facilitative (positive) or when the number of negative (competitive/exploitative) interactions is an even number. The loop has a negative feedback property when the number of negative interactions in the loop is an odd number. In fact, this last rule applies to feedback loops of any length, with an odd number of negative interactions being stabilizing, and an even number of negative interactions being destabilizing.

### Appendix 2: Environmental data acquisition and modelling

Meteorological data for the study site and period (1985-2024) were obtained from the European gridded observational dataset E-OBS version 31.0e (2), available at the 0.1°×0.1° grid. Daily precipitation and mean, minimum, and maximum air temperature were extracted for the grid cell corresponding to the study location and then aggregated to monthly values. The meteorological forcing data were used to drive the mesoscale Hydrologic Model [mHM (3, 4)], in order to derive basic hydrological variables of the study domain. mHM is a spatially-explicit distributed hydrologic model based on the multiscale parameter regionalisation concept. From existing European mHM simulations, we extracted daily site-level time series of modelled snowmelt and upper-layer soil moisture, representing approximately the upper 30 cm of the soil profile. Snowmelt was aggregated to monthly totals and soil moisture to monthly means. These variables were used as model-based indicators of local hydroclimatic conditions at the study site. Reconstructed time series were consistent with the point-scale framework (5), they also captured well-known recent European drought periods, including 2018-2020 and 2022, providing a qualitative check on the hydroclimatic variability represented by the simulations (6, 7).

Vegetation dynamics were characterised using NDVI derived from NOAA Climate Data Record products in Google Earth Engine. Daily NDVI has been extracted from the AVHRR NDVI Version 5 product (8), and the VIIRS NDVI Version 1 product (8), both products provide daily NDVI on a 0.05°×0.05° global grid. Daily values extracted at the study site were converted using the standard scale factor and aggregated to monthly values.

Long-term atmospheric pollution (1985–2024) was assessed using concentrations of suspended particulate matter — SPM (µg/m$^3$). Data were obtained from the monitoring network of the Czech Hydrometeorological Institute (CHMI) and their complementary regional stations. Monthly mean concentrations were calculated from average daily measurements. Because no single monitoring station covered the entire study period, the time series was compiled from four stations located in the vicinity of the study area: Horní Jiřetín (7 km from the study area; 19865–1995), Fláje (3 km; 1995–2002), Zinnwald (19 km; 1996–2000), and Rudolice v Horách (11 km; 2001 and 2003–2024). Inter-station distances of these stations ranged from 9 to 43 km. Measurements of SPM were available for the period 1985–2003. From 2004 onwards, when only $PM_{10}$ concentrations were routinely monitored, $PM_{10}$ values were converted to SPM equivalents using a standard conversion coefficient of 1.2 [Council Directive 1999/30/EC of 22 April 1999. Official Journal L 163, 29/06/1999 P.0041-0060; (9)].

### Appendix 3: Multivariate state-space Bayesian model (MDGAM)

Observed counts of trapped animals $Y_{s,t}$ for series of species $s$ at time $t$ were assumed to follow a negative binomial distribution to account for overdispersion, with $Y_{g,t} \sim NegBin(\mu_{s,t}, \Phi)$, where $\mu_{s,t}$is the expected abundance and $\Phi$ is the overdispersion parameter. The observed abundances were modelled via a log link function, as follow:

$$\log(\mu_{s,t}) = \ series_s + \sum_{l=0}^{4} f_{l,g[s]}\,(SPM_{t-l}) + z_{s,t}$$

With $series_s$ being a parametric intercept for each time series, and $SPM_{t-l}$ the lags $l$ of atmospheric pollution (from 0 to 4 months). The term $f_{l,g[s]}$ is a hierarchical random-effect smooth (modelled as bs = 're' in "*mvgam*" R package) that estimated a specific Beta slope coefficient for each ecological guild $g$ at lag $l$, where species $s$ belongs to guild $g$. The latent temporal dynamics were captured by a state space component $z_{s,t}$, representing the series-specific trend (one for each species). The dynamics of these latent trends were modelled using a vector autoregressive process of order one (AR1), defined in the latent trend formula as:

$$z_t = f_{global}\ t\ + f_{dev,s}\ t\ + \boldsymbol{A} z_{t-1} + \varepsilon_t$$

Where $z_t$ is the vector of latent states across all series at time *t*. The deterministic component of the trend includes a shared global smooth for time $f_{global}$ (a cubic regression spline, modelled as bs = 'cr' in "*mvgam*" R package), and series-specific smooth deviations from that global shared latent trend $f_{dev,s}$ (implemented with a factor –interaction spline with a null-space penalty, noted as bs = 'sz', xt = list(bs = 'cr'). The stochastic component is driven by the transition matrix $\boldsymbol{A}$ which captures temporal autocorrelation and cross-series dependencies, and $\varepsilon_t$ the multivariate normal error term with a correlated covariance matrix.

The model was implemented in a Bayesian framework using the "*mvgam*" R package in R, using *CmdStanR* for Hamiltonian Monte Carlo sampling. To ensure robust posterior convergence and avoid sampling issues, a non-centered parametrization was used, and initial values were set to zero. We ran the model using default priors in "*mvgam*". Convergence was checked using standard diagnostic of chain mixing (Rhat close to one and adequate effective sample sizes).

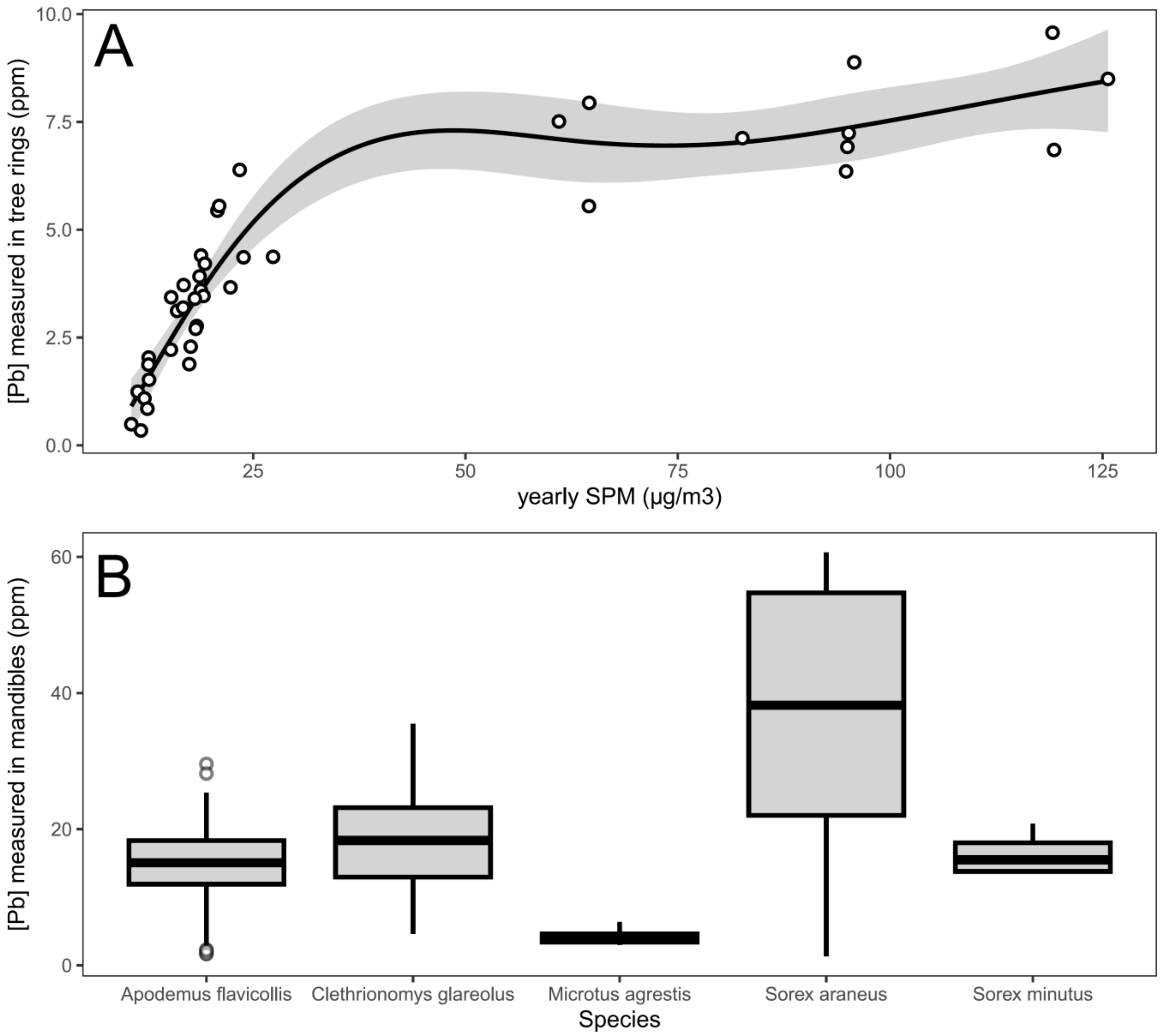


**Fig. S1**: (A) Relationship between measured yearly lead (Pb) in tree rings on the study site and yearly averaged SPM levels used as an index of pollution (GAM, $R^2$=0.87). Tree rings have been sampled in Norway spruces in 2025 on the Ore mountain plateau, near the study plots. (B) Boxplots showing marked differences of lead (Pb) concentrations in mandibles among species (122 mandibles sampled in 2017), with very small and poorly variable concentrations in the strict herbivore *Microtus agrestis*, intermediate concentrations in the omnivores *Apodemus flavicollis* and *Clethrionomys glareolus*, and relatively higher concentrations in the insectivore *Sorex araneus*. Lead in each tree ring and each mandible has been quantified using Inductively Coupled Plasma – Mass spectrometry (ICP-MS) in the geoscience laboratory of Prague (CZU, FZP), following routine protocols; e.g., Chrastný et al. (11).

### Interpretation of measured Lead concentrations

The concentrations differed among species within a guild, with *Sorex minutus* showing smaller concentrations than *Sorex araneus*, the former showing concentrations that were comparable to the ones of the omnivores. This could be explained as insectivores feed on several trophic levels in the invertebrate food webs, and as *Sorex minutus* is significantly smaller in size than *Sorex araneus,* thus probably feeding at a lower trophic level within insects/invertebrates. Therefore, we hypothesize that concentrations of lead in mandibles exhibit guild- and species-specific trophic behavior (or diets) as guilds and species feed at different trophic levels (with grass being probably little contaminated compared to seeds, lichens, and insects consumed by omnivores, which is overall relatively less contaminated than a diet composed essentially of insects by insectivores, which themselves vary in lead concentrations, depending on the invertebrate trophic levels).

Overall, with a grand mean average of lead in mandibles over 20 ppm in 2017 (Fig. S1B), and such a saturating positive relationship between SPM and lead in tree rings (Fig. S1A), we argue that the effects of pollution at the peak and until the end of the polluted period were toxic to small mammals, altering their physiology, behavior and demographics, depending on the sensitivity to pollution exhibited by each species (and we warn our readers that exposure does not equate to sensitivity to pollution, even if we believe that both should be broadly linked positively). Second, these values indicate that lead exposure is still substantial now, but could have strongly reduced after that the pollution peaked. As a comparison, small mammal individuals sampled in the UK in 1978 in control and smelting/mine contaminated sites, showed concentrations similar to ours (around 10/20ppm) in control sites, while concentrations were >100/ 200 ppm in contaminated sites (10). Please note that Lead (Pb) is non-essential and (hyper) toxic to animals even at low concentrations.

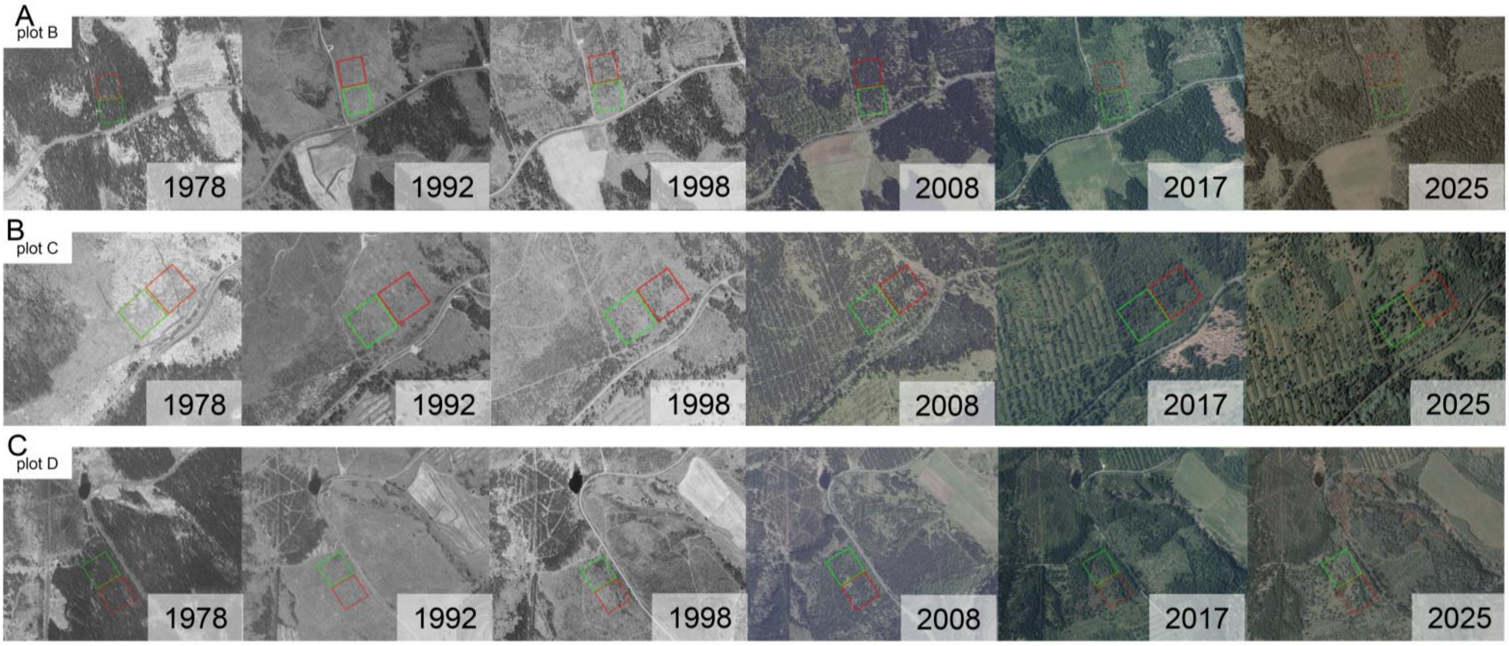


**Fig. S2**: Temporal series of orthophotos showing the changes in land use in our study plots. In 1986, the trapping plots were situated in clear-cuts created after logging. These areas were subsequently planted with predominantly non-native blue spruce seedlings. Compared to native northern Norway spruce forests, Colorado blue spruce stands do not develop mature, dense canopies. Indeed, even after 20–30 years of growth, blue spruce stands usually remained below 10 m in height and formed open areas dominated by hairy reed grass (where each plot was located) among discontinuous spruce stands. The area was subjected to minimal forestry intervention until Colorado blue spruce stands were removed and replaced with native Norway spruce plantings. This occurred sequentially in autumn 2012 in plot B, in spring 2017 in plot C, and in autumn 2018 in plot D. The 1-ha plots used for small-mammal trapping are shown as coloured squares: in green for spring plots, and in red for autumn plots.

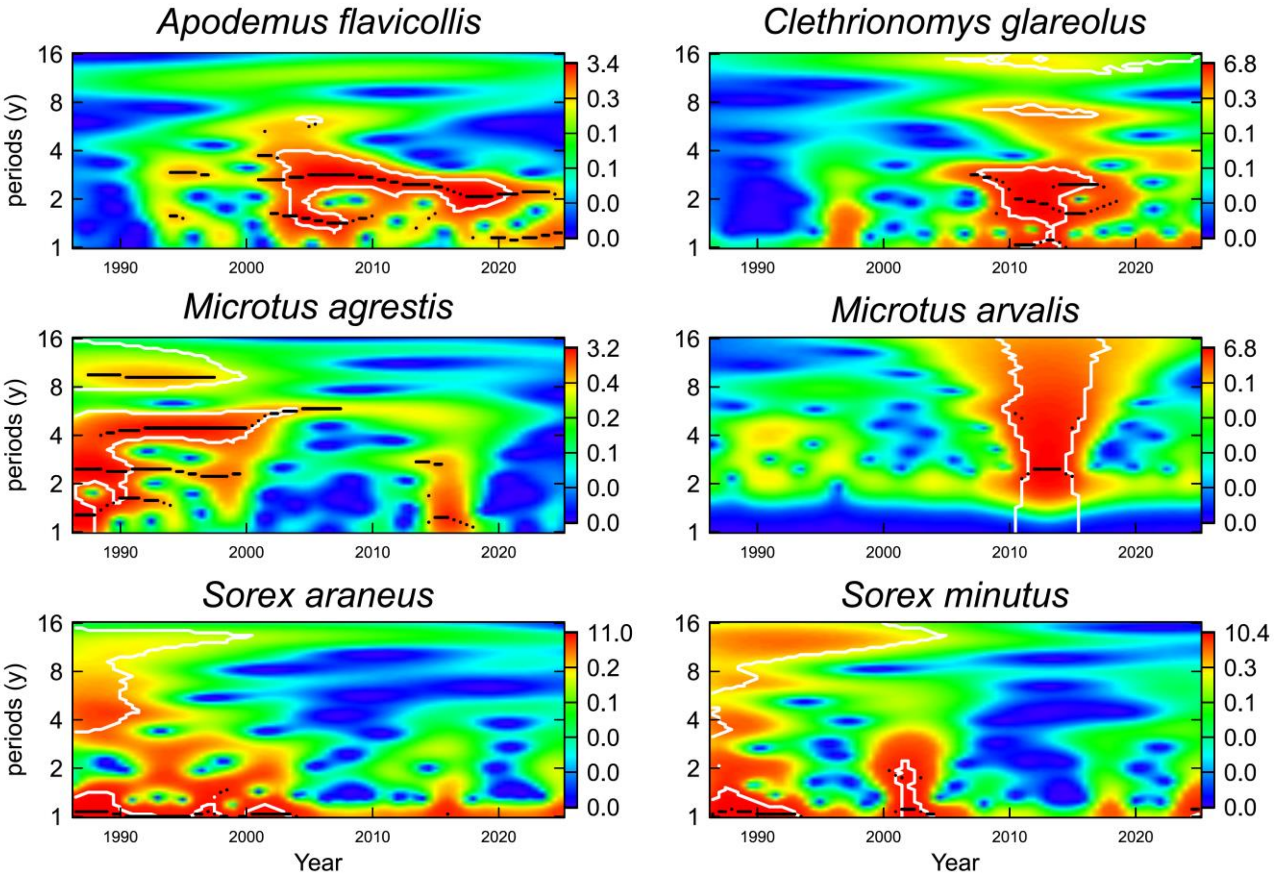


**Fig. S3**: Wavelet power spectrums (WPS) of small mammal population dynamics.

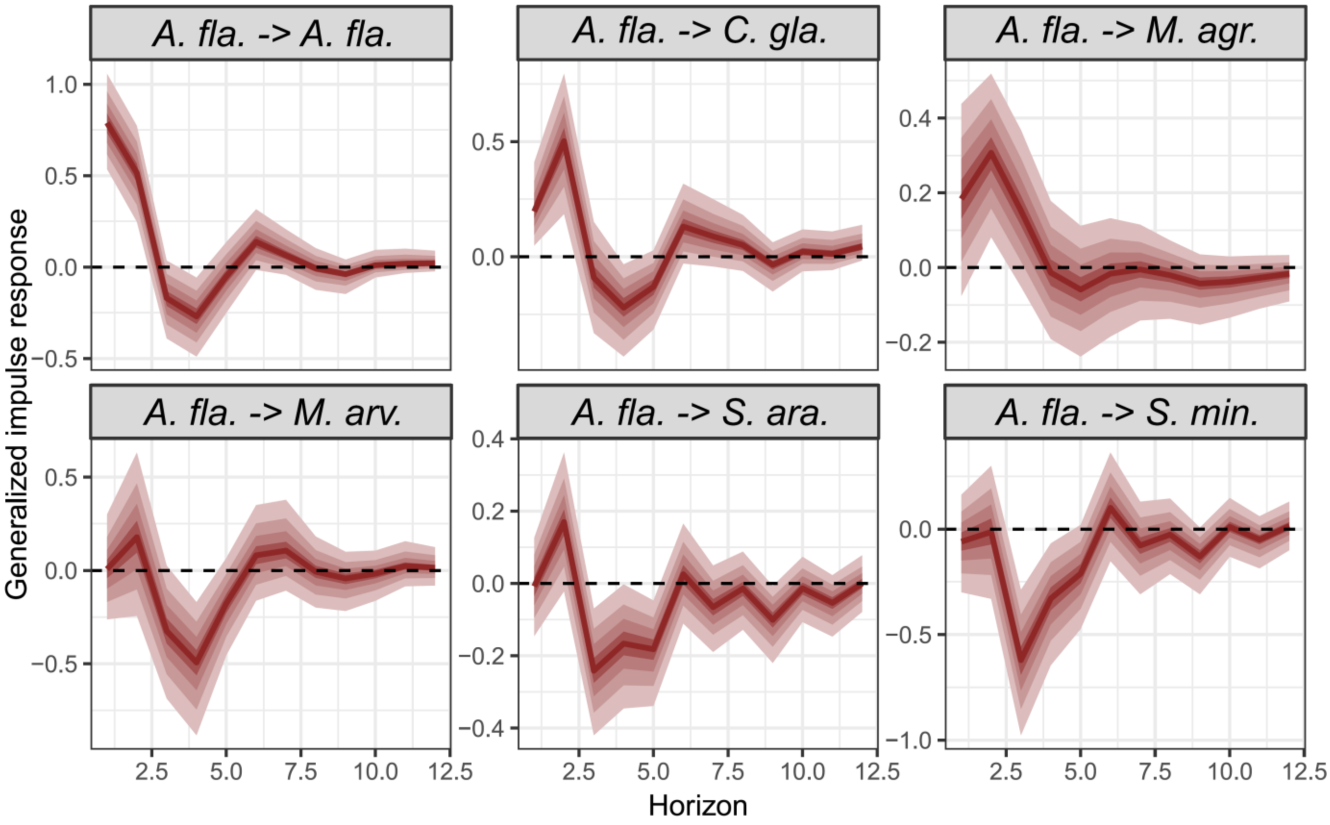


**Fig. S4**: Impulse response functions to shock increases of *Apodemus flavicollis* on its own population and on each other species are presented with 90%, 60%, 40%, and 20% credible intervals along forecast horizons (up to 12.5 time steps, equivalent to 6 years).

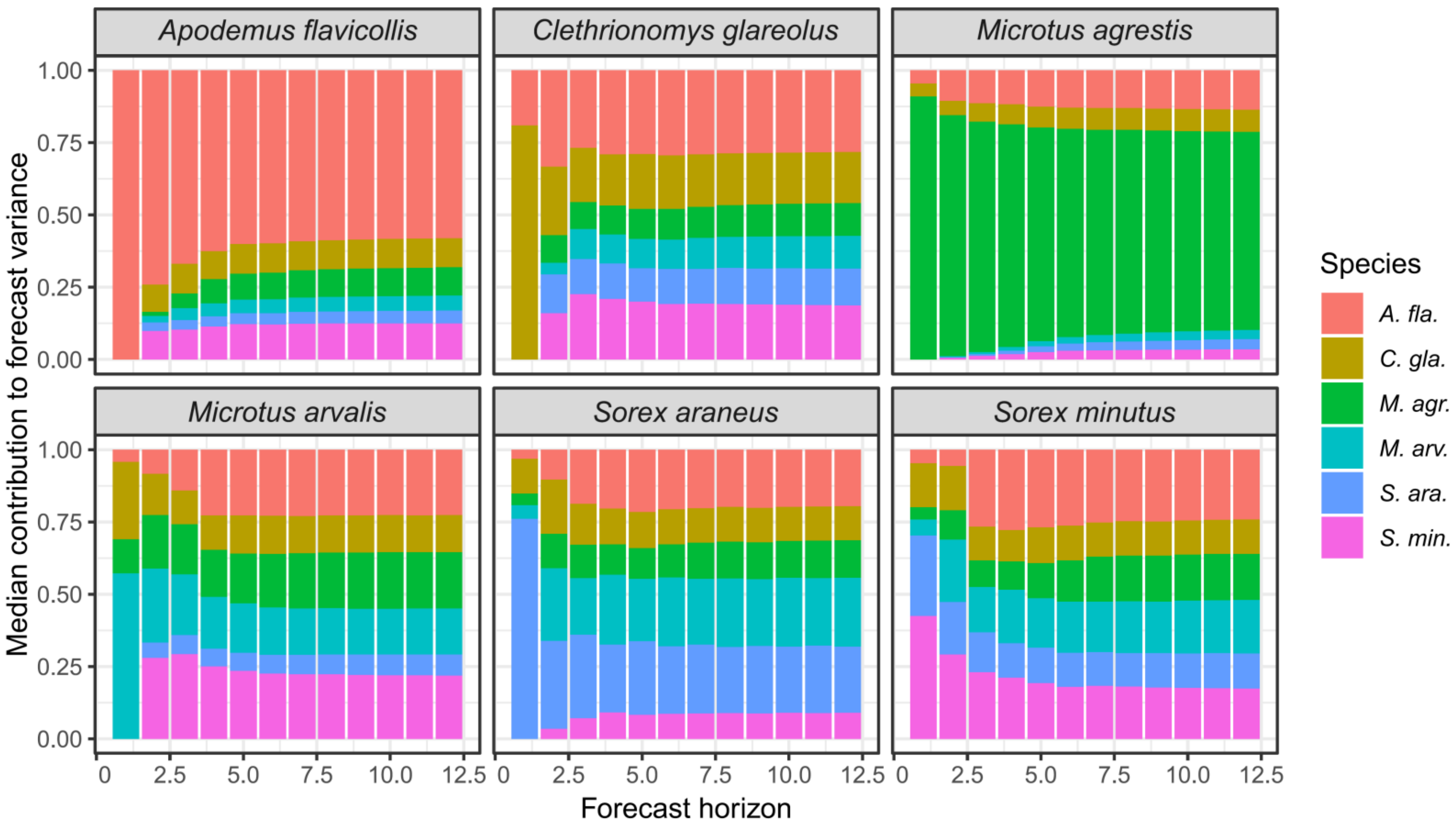


**Fig. S5**: Decomposition of forecast variance to generalized impulse functions.

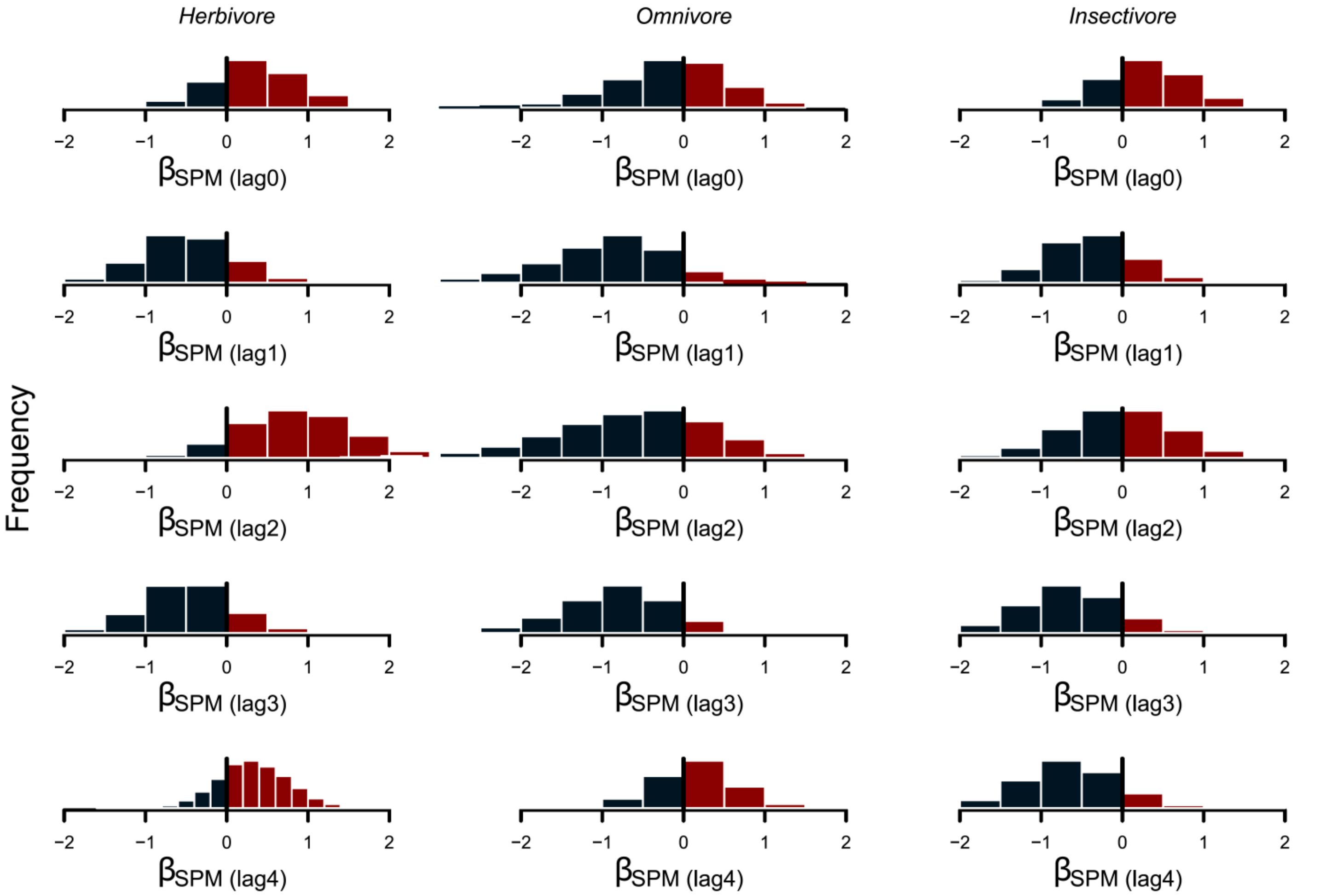


**Fig. S6**: Lagged effects of atmospheric pollution (SPM levels) by guild. Histograms show the posterior distributions of beta coefficients for the lagged linear effects of SPM increases (lags 0–4 months) for each guild. Black bars indicate the negative part, and the red bars, the positive parts of the posterior distributions of beta estimates.

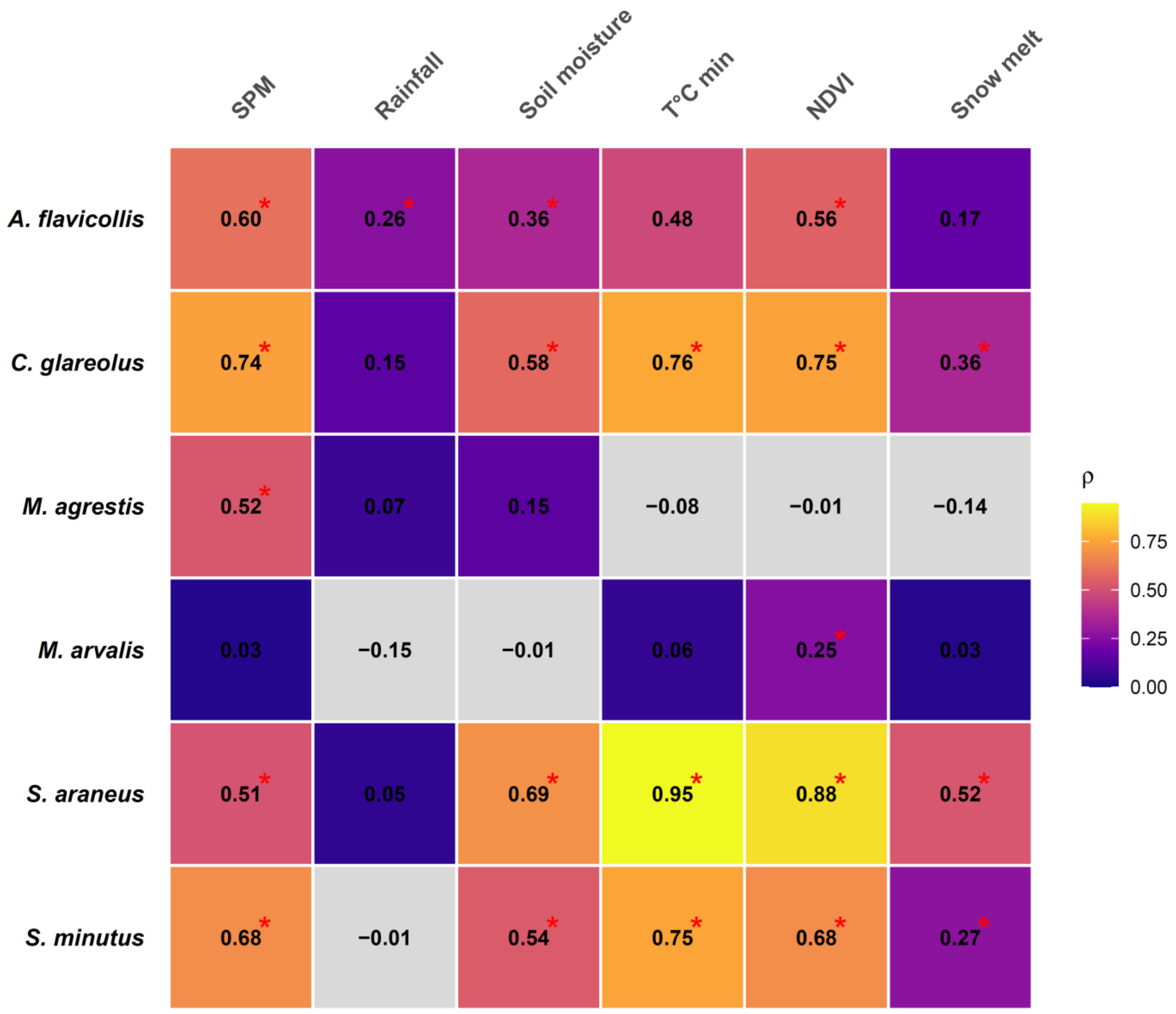


**Fig. S7**: CCM of the causal effects of environmental drivers on each species (cross skills *ρ*) represented as a heatmap. Red asterisks show *p* values < 0.05 against a null distribution of 1000 seasonal surrogates.

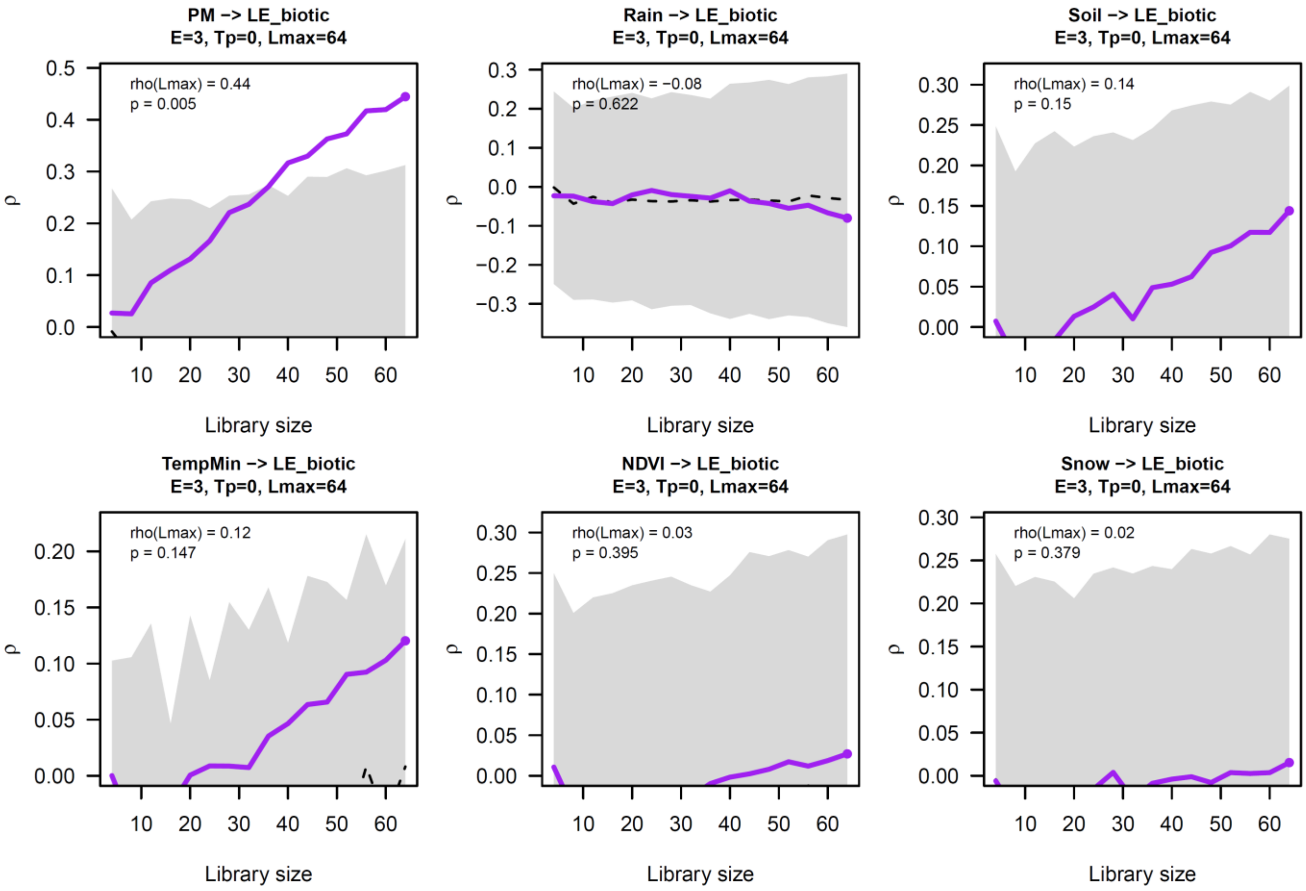


**Fig. S8**: Convergent-cross mapping (CCM) plots showing the convergence, or lack thereof, of observed prediction skills (purple line) as library size increases for each significant environmental driver → local lyapunov exponents "LLE_biotic" (dynamical instability or chaos). "PM" stands for suspended particulate matter (SPM), "Soil" for soil moisture, and "Snow" for snow melt. Grey bands show the 95% null distribution based on 1000 seasonal surrogates.

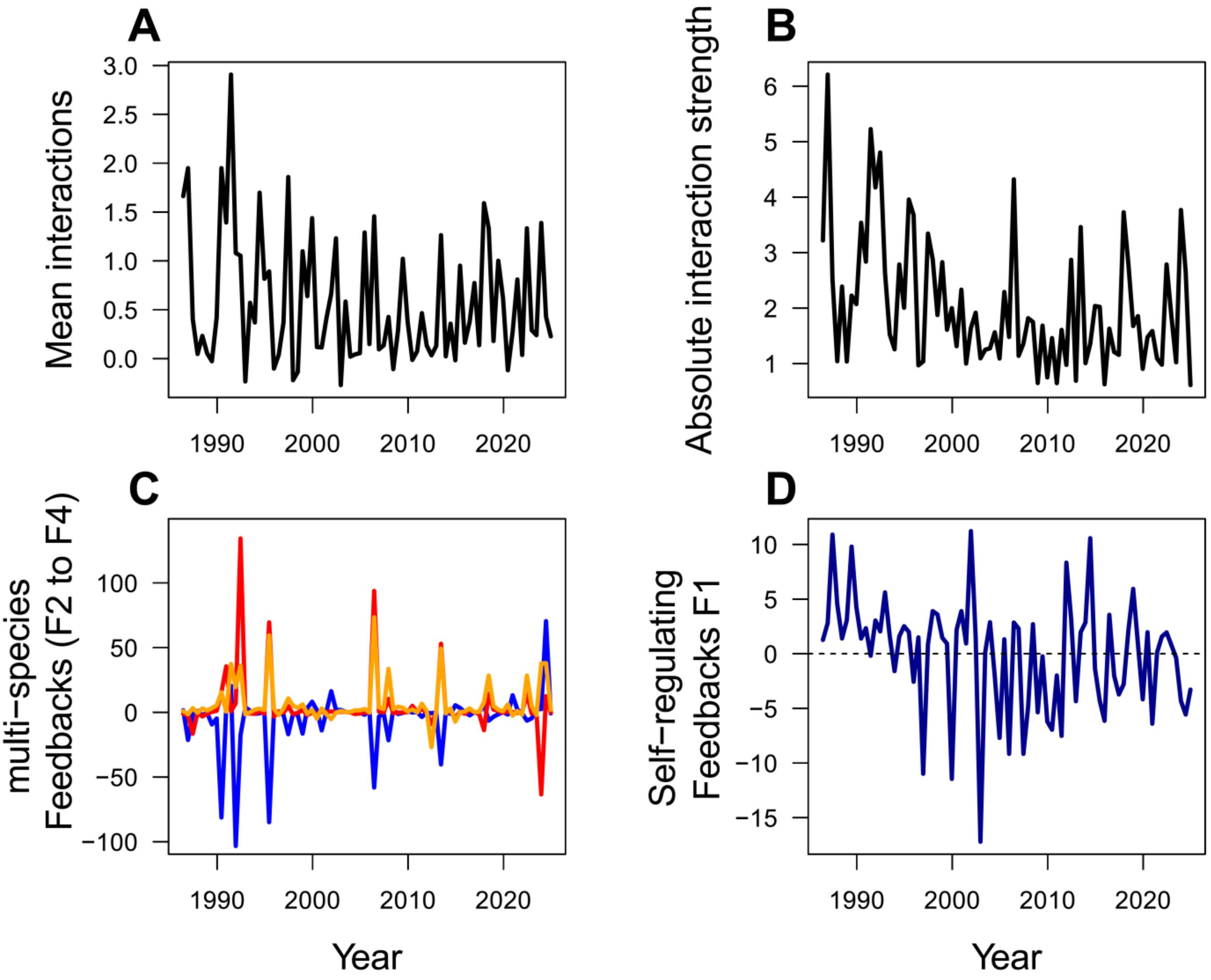


**Fig. S9a**: Time varying interaction metrics, showing the time series of the average of signed interactions (A), the average of absolute interactions (B), the feedbacks "F" of the two, three, and four species loops (F2: orange, F3: red, and F4: blue, respectively) (C), and the average of the self-regulating feedbacks F1 of the one species loops in dark blue (D).

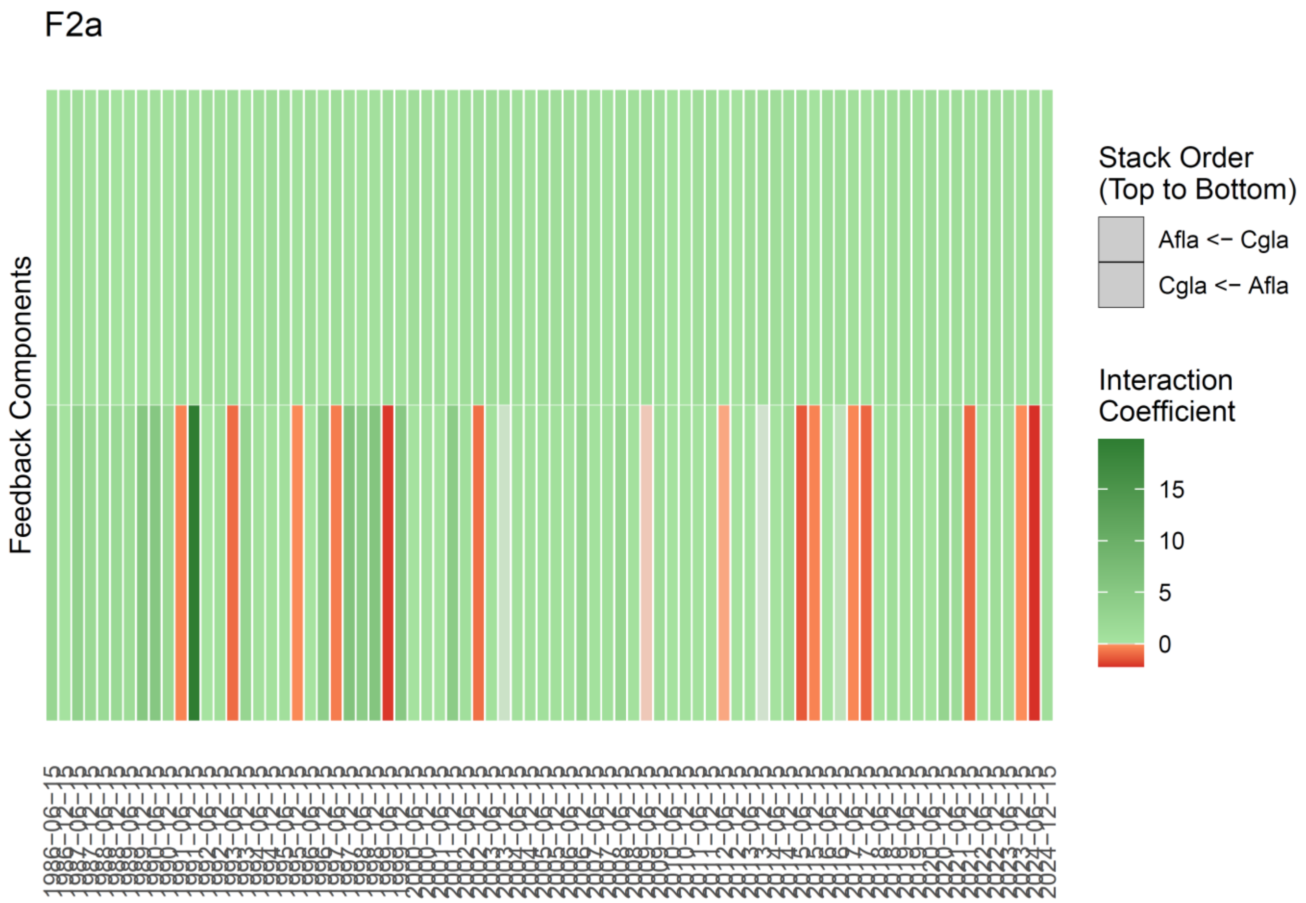


**Fig. S9b(i)**: Interaction coefficients between each species in the two-species feedback loops F2a, showing an increase of competitive interactions of *A. fla.* on *C. gla.* as the polluted period came to an end. From 2002 to 2011, the loop became facilitative to show again shocks of competitive interactions from *A. fla.* in the last decade. This loop showed negative spikes at these two moments but was otherwise destabilizing.

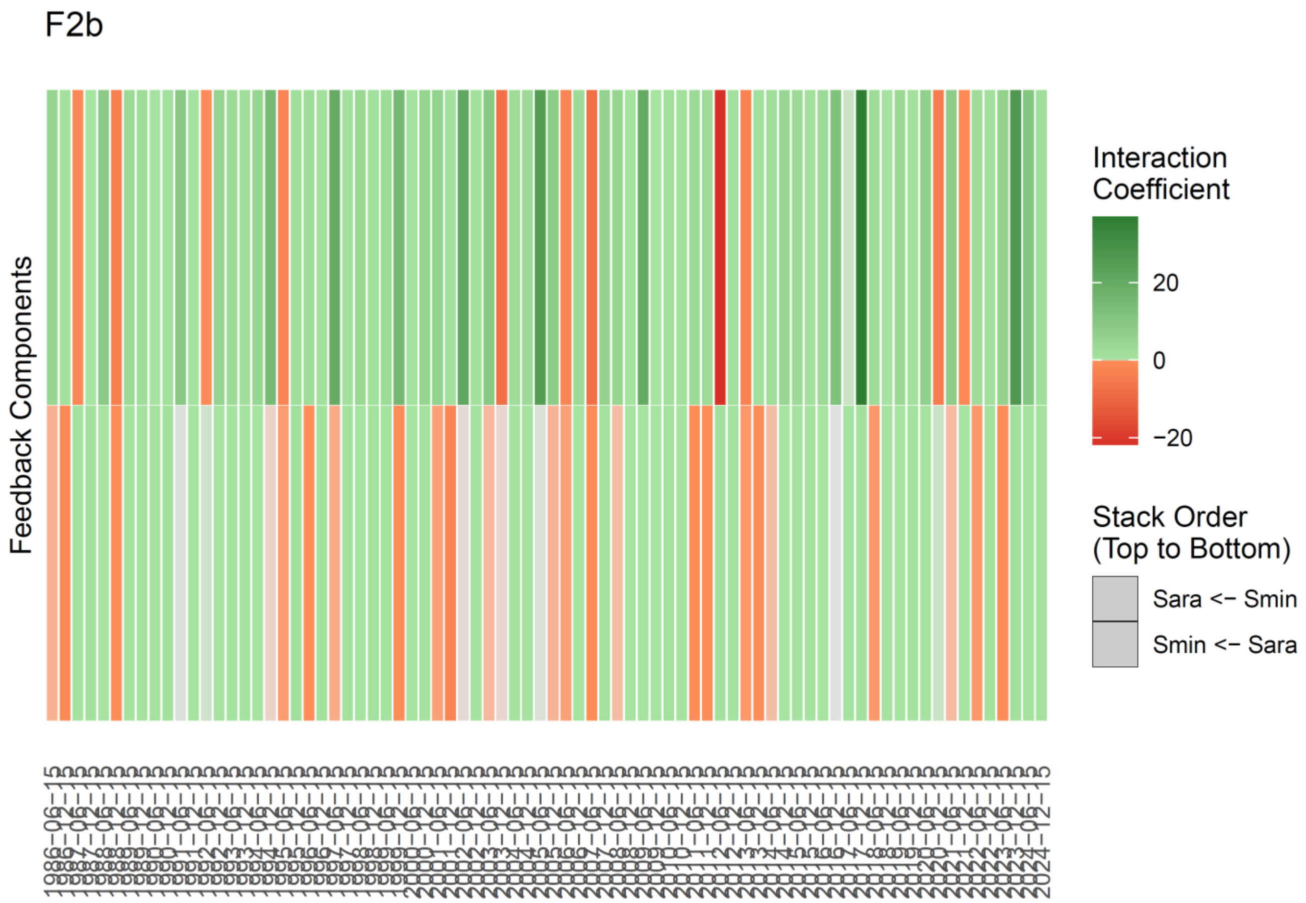


**Fig. S9b(ii)**: Interaction coefficients between each species in the two-species feedback loops F2b. The couple of shrews do not see a marked evolution, with overall either facilitative or two competitive interactions but a marked negative spike from *S.min* to *S. ara.* in spring 2012. This loop is overall destabilizing, but show several negative effects.

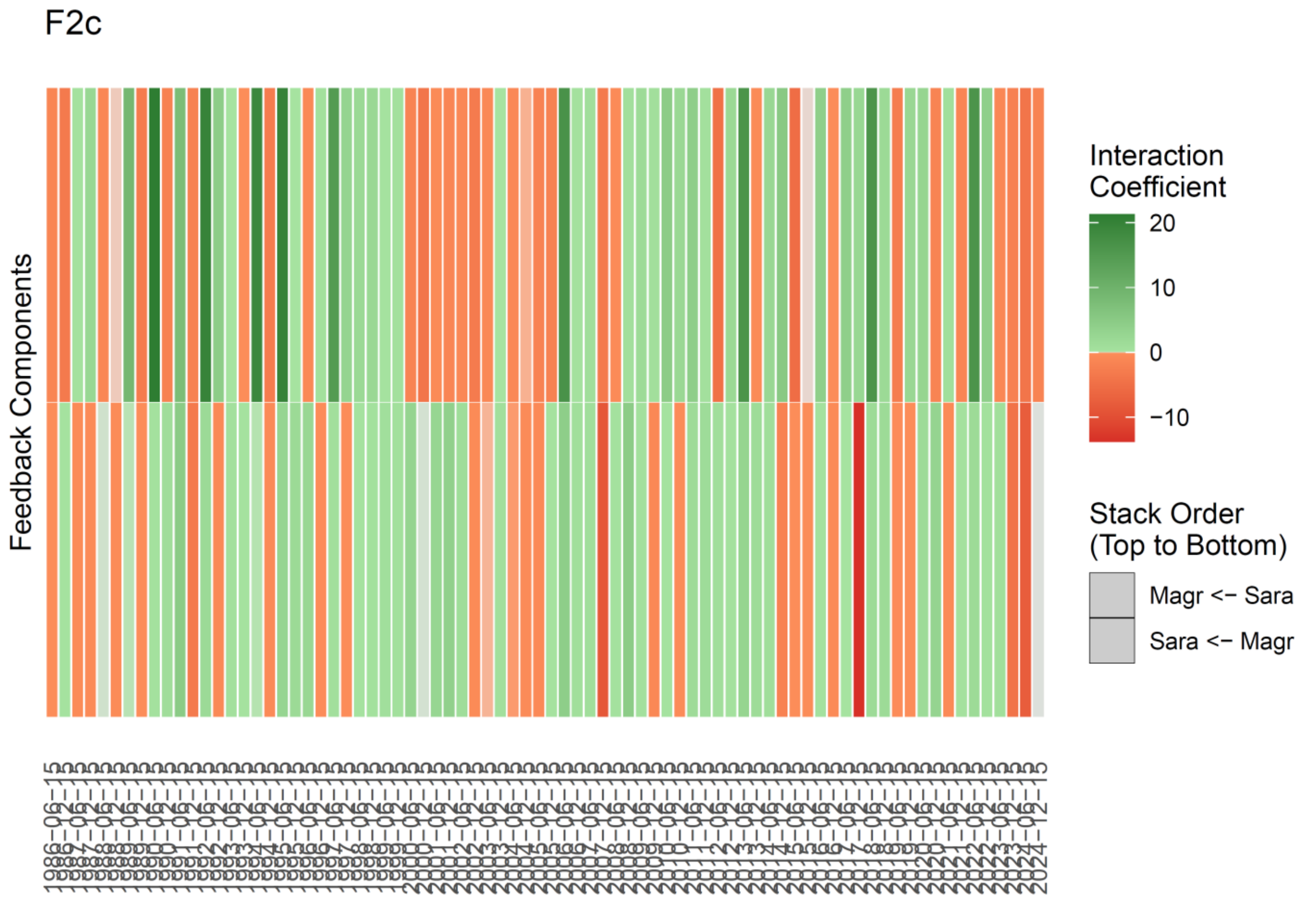


**Fig. S9b(iii)**: Interaction coefficients between each species in the two-species feedback loops F2c. Interactions between *S. ara.* and *M. agr.* shows an alternation between competition and strong facilitation during the polluted period, and after 2 years of facilitative interactions (1998 and 1999), the loop showed a block of competition during 2.5 years (2000-2002), before to start alternating again between competition and facilitation.

F3a

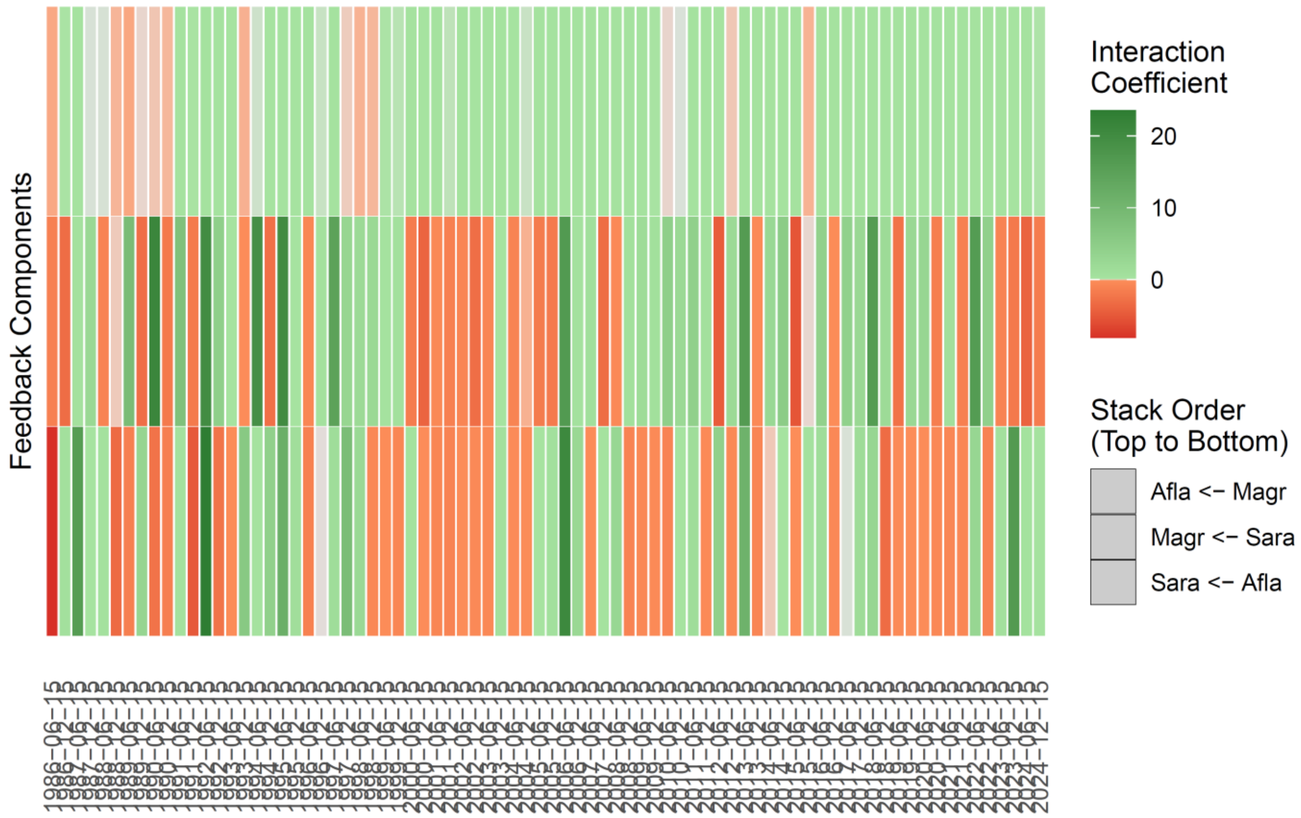


**Fig. S9b(iv)**: Interaction coefficients between each species in the three species feedback loop F3a, showing in the first decade (1986-1999) a regime marked by a few strong competitive interactions (red) and several strongly facilitative (green) interactions that we interpret as overcompensatory feedbacks (apparent facilitation). The decade 2000-2003 shows a block of two competitive interactions of *A. fla.* on *S. ara.* and from *S. ara.* on *M. agr.*, with positive interactions for *M. agr.* on *A. fla.*, which are destabilizing conditions (just before that we identified marked signed of a critical transition with a Bayesian change-point model and with multivariate EWS statistics; see main text). The last 2/3 of the study shows a mixed pattern of competitive vs facilitative interactions.

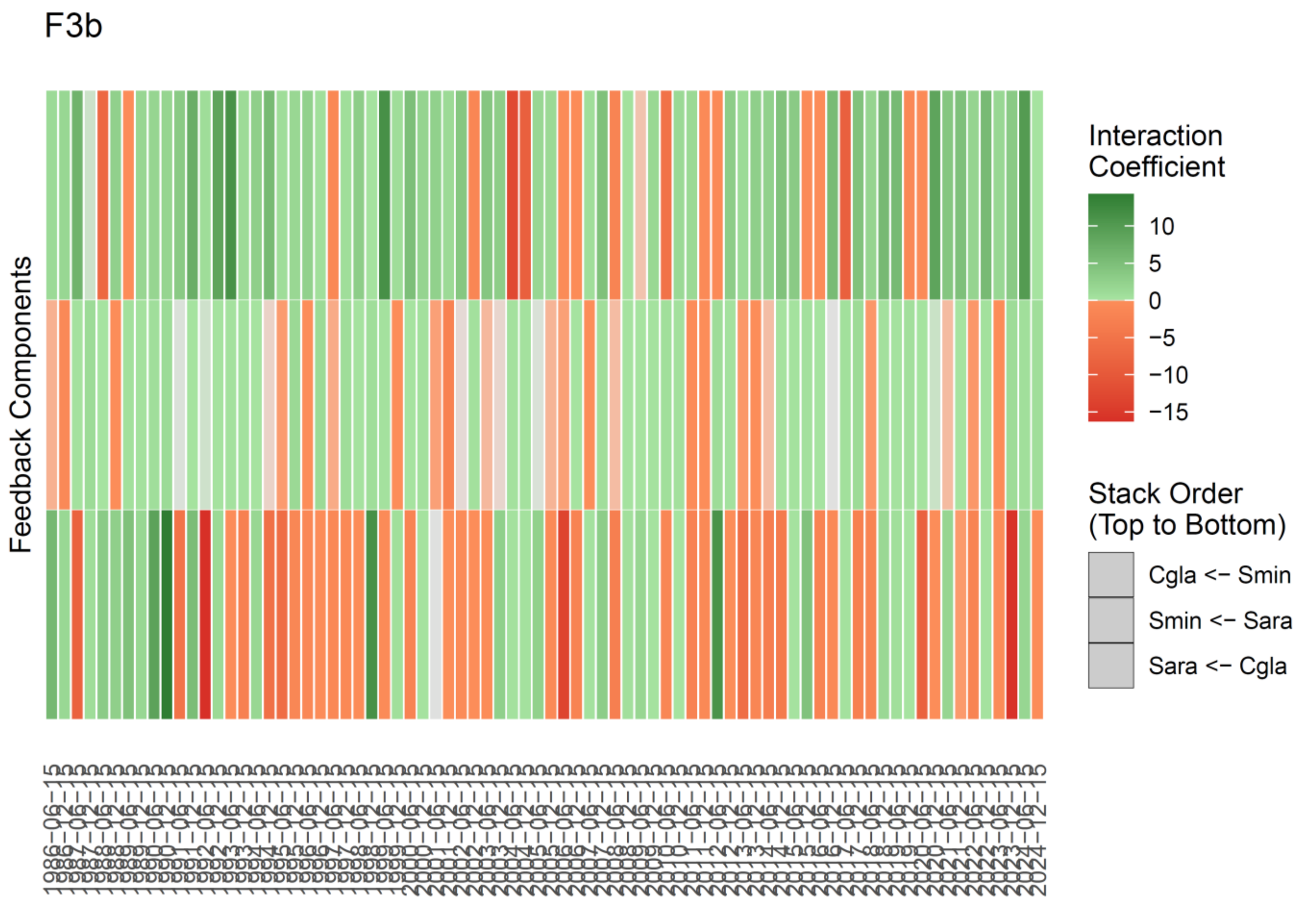


**Fig. S9b(v)**: Interaction coefficients between each species in the three species feedback loop F3b. The interactions from *S. minutus* to *C. glareolus* were positive up to ~2000, and showed then marked negative spikes in 2004, just before the critical transition was detected. *C. glareolus* has overall negative effects on *S. araneus*. Otherwise, even numbers of negative interactions were common in this feedback loop, suggesting its destabilizing (positive) nature on system’s dynamics.

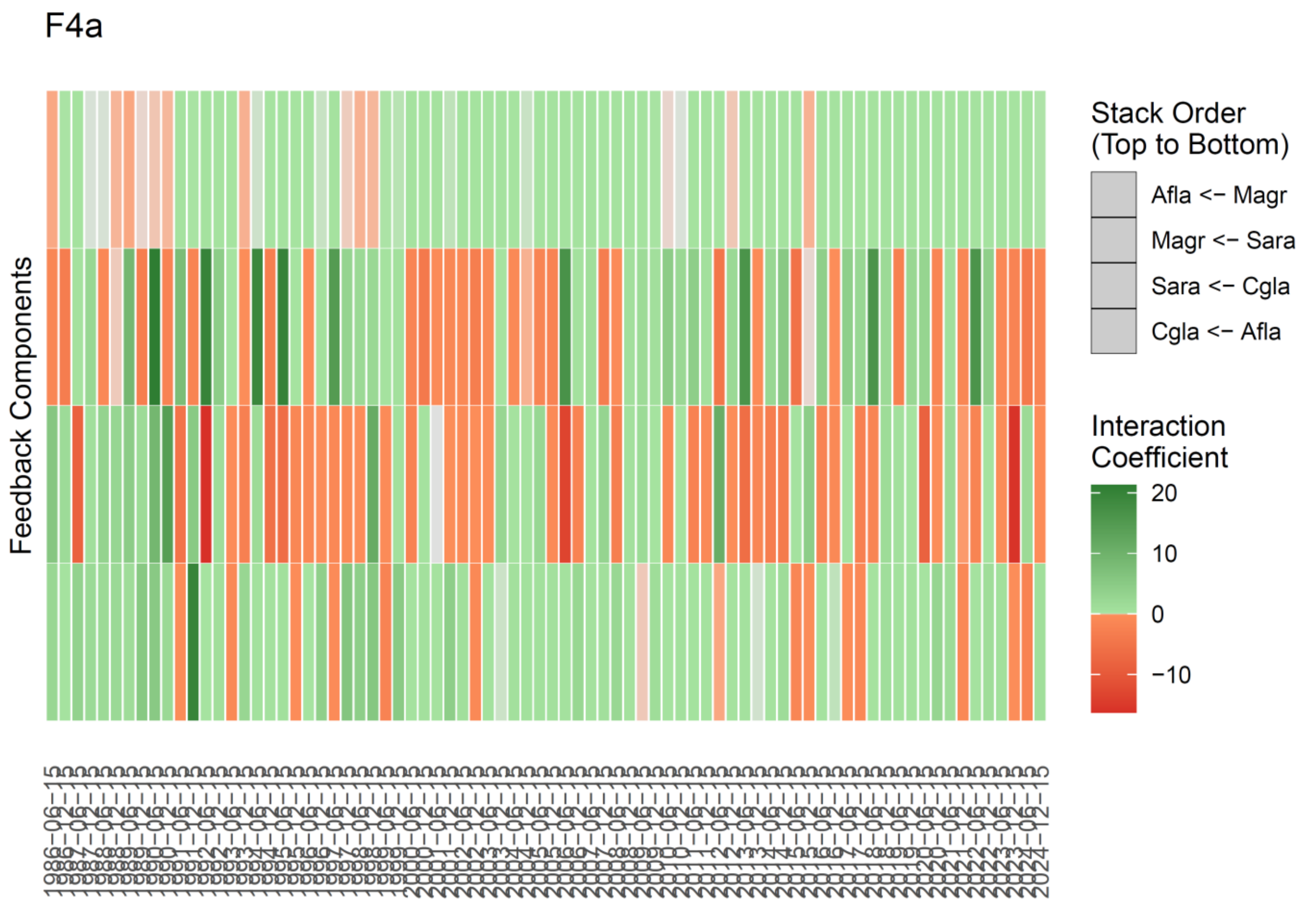


**Fig. S9b(vi)**: Interaction coefficients between each species in the four species feedback loop F4a. This four-species feedback loop shows in the first decade (1986-1997) a regime marked by a few strong competitive interactions (red) and several strongly facilitative (green) interactions that we interpret as overcompensatory feedbacks (apparent facilitation). The presence of odd numbers of negative interactions contributes to this pattern characterized by strong interactions strengths and abrupt negative products of interactions in the feedback loop. In 2000-2002, we observed a period of weakly positive interactions, before a mixed pattern until the end of the study.

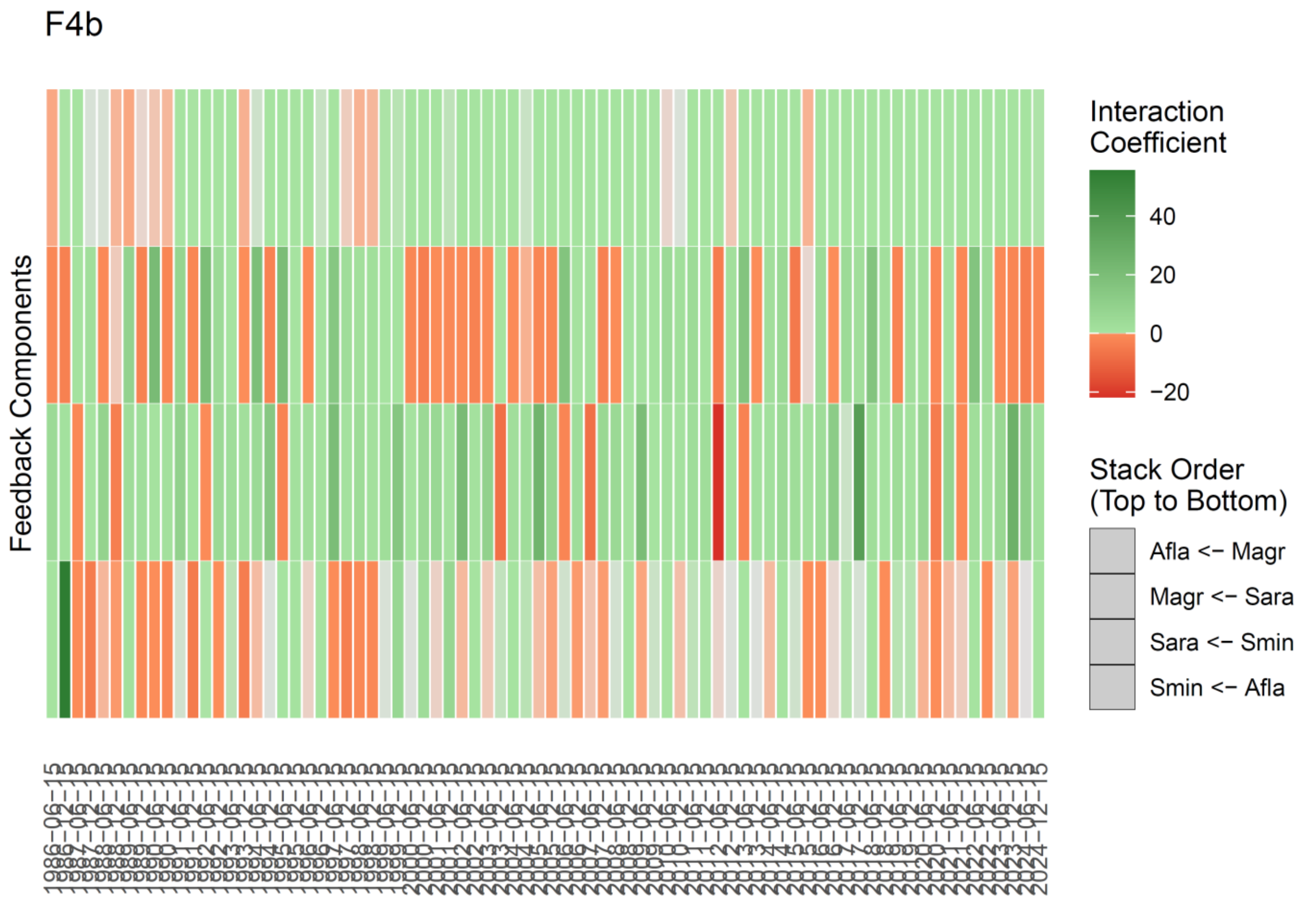


**Fig. S9b(vii)**: Interaction coefficients between each species in the three species feedback loop F4b. The loop F4b shows a similar pattern than F4a, with noticeably stronger negative interactions from *A. fla.* on *S. min* during the polluted era. Both loops tended to show more positive products in the last decade.

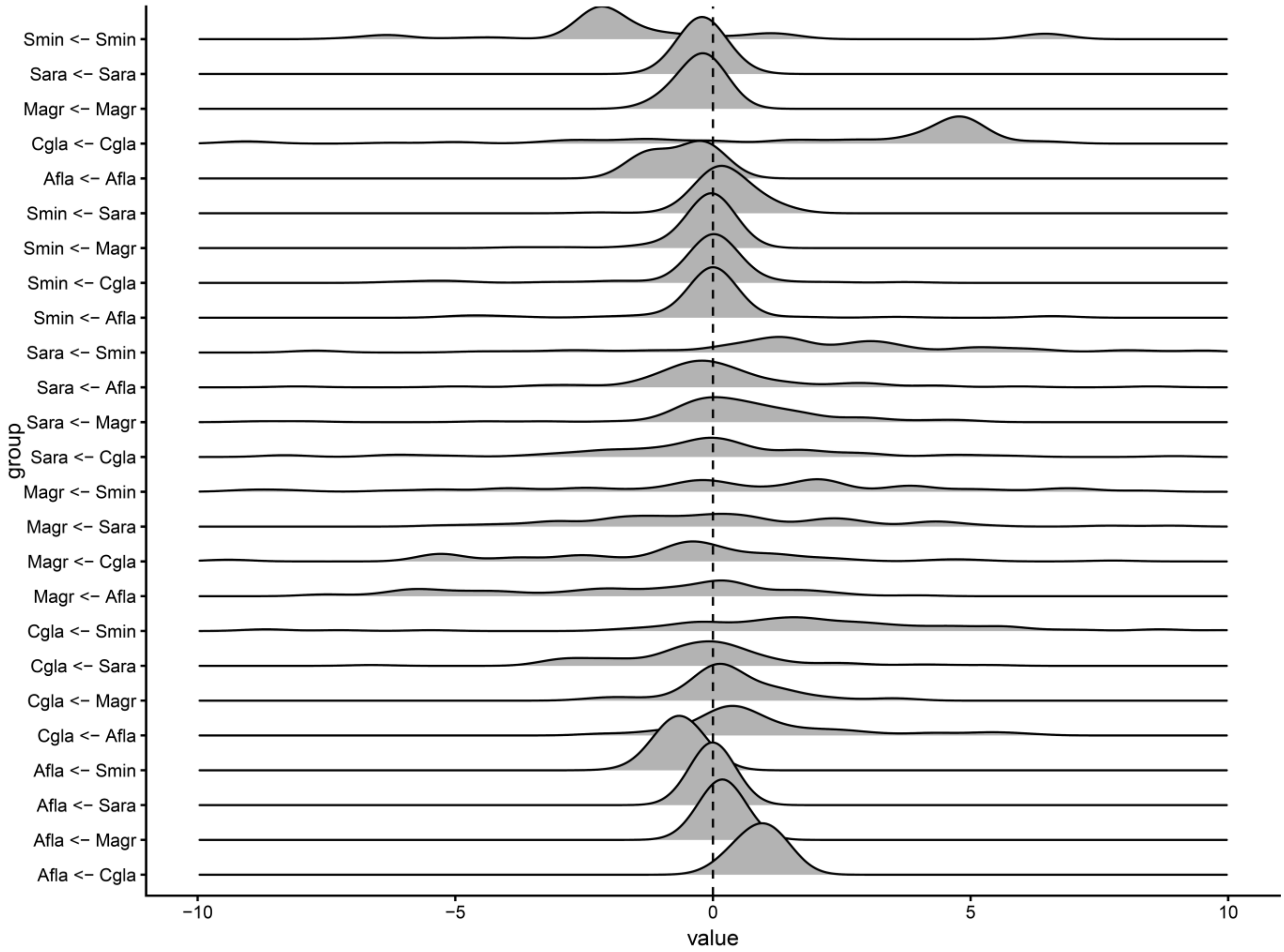


**Fig. S10**: Distributions of estimated time-varying interaction coefficients between each of the twenty interspecific pairs and five intraspecific pairs (self-regulation of self-facilitation) in the network from multivariate S-maps. Negative values are competitive interactions, and positive values are facilitative interactions.

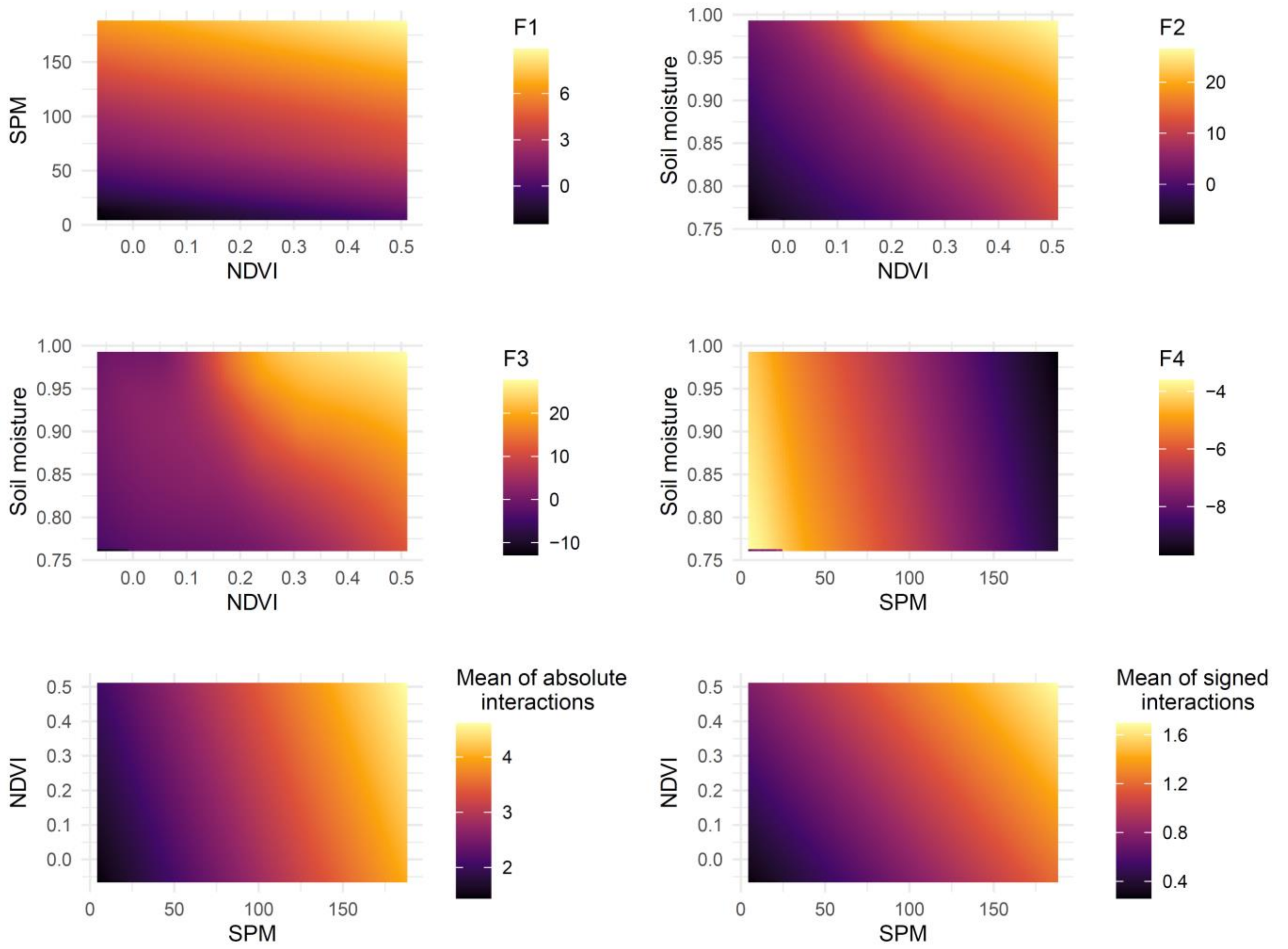


**Fig. S11**: Surface plots of how chosen environmental drivers (based on their CCM predictive skills and p-values in Fig. 2E, main text) affect the interaction metrics (100 iterations for each S-map).

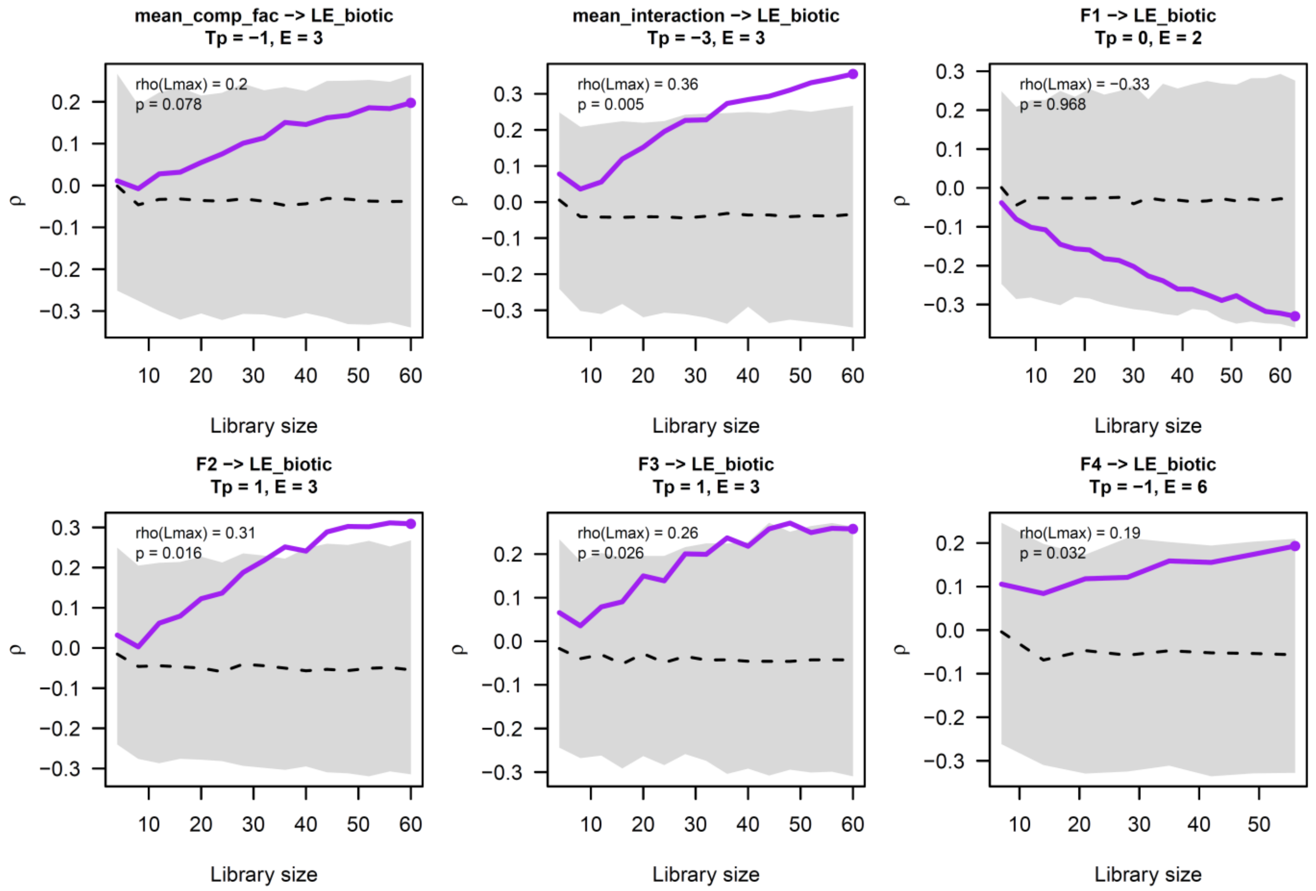


**Fig. S12**: CCM convergence plots of the cross mappings between the metrics of the interaction network and the local Lyapunov exponents (LLEs, or chaos). The purple line show the convergence of observed rho values as the library length increases. “mean_comp_fac” stands for mean signed interactions, “mean_interaction” for mean absolute interactions, and “LE_biotic” for local lyapunov exponents (LLEs). The shaded grey bandwidth shows the distribution of a 1000 null seasonal surrogates.

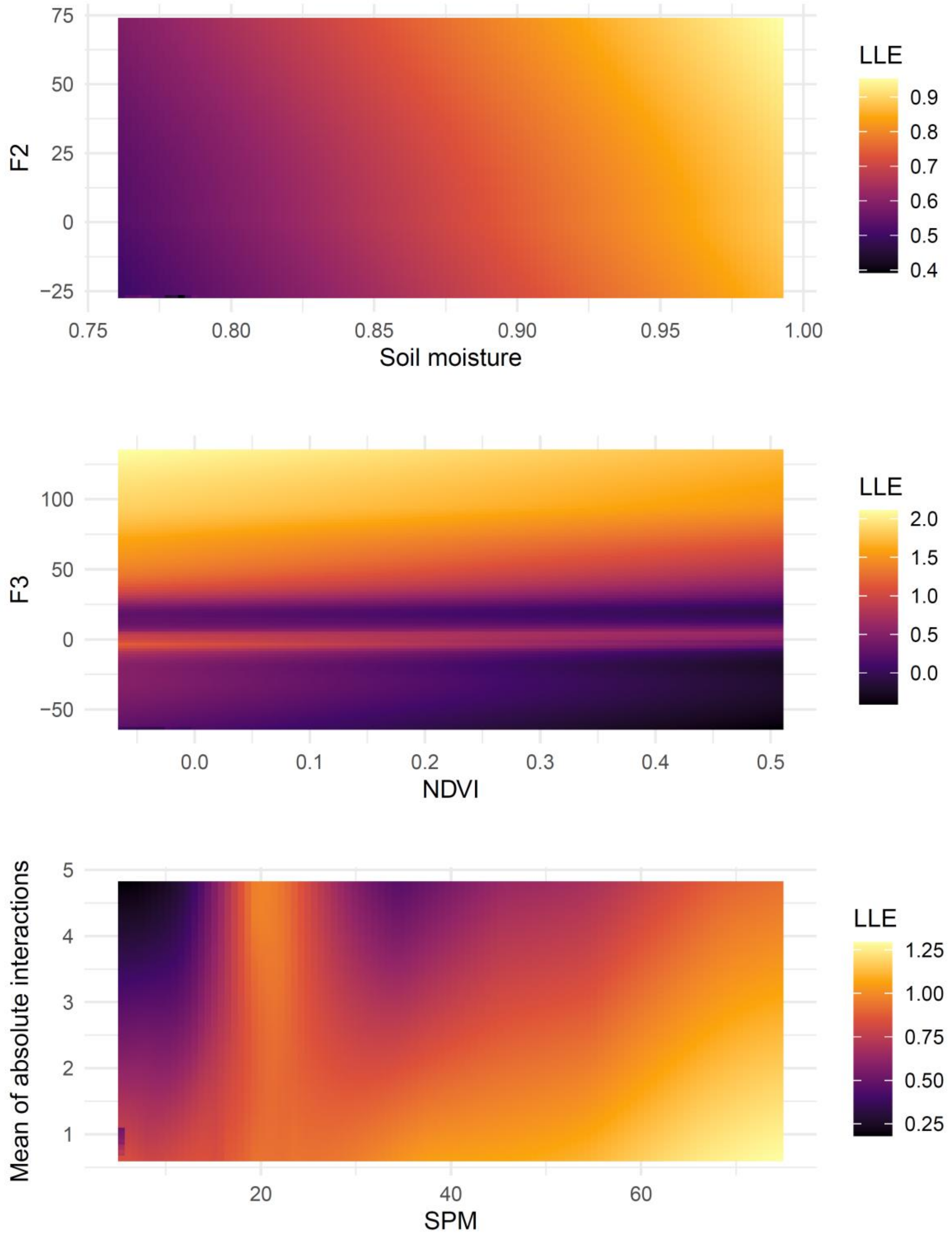


**Fig. S13**: Surface plots of how interaction metrics (the three ones cross mapping with local lyapunov exponents, LLE) affect jointly with the main environmental drivers affecting those same interaction metrics, the LLEs (100 iterations for each S-map).

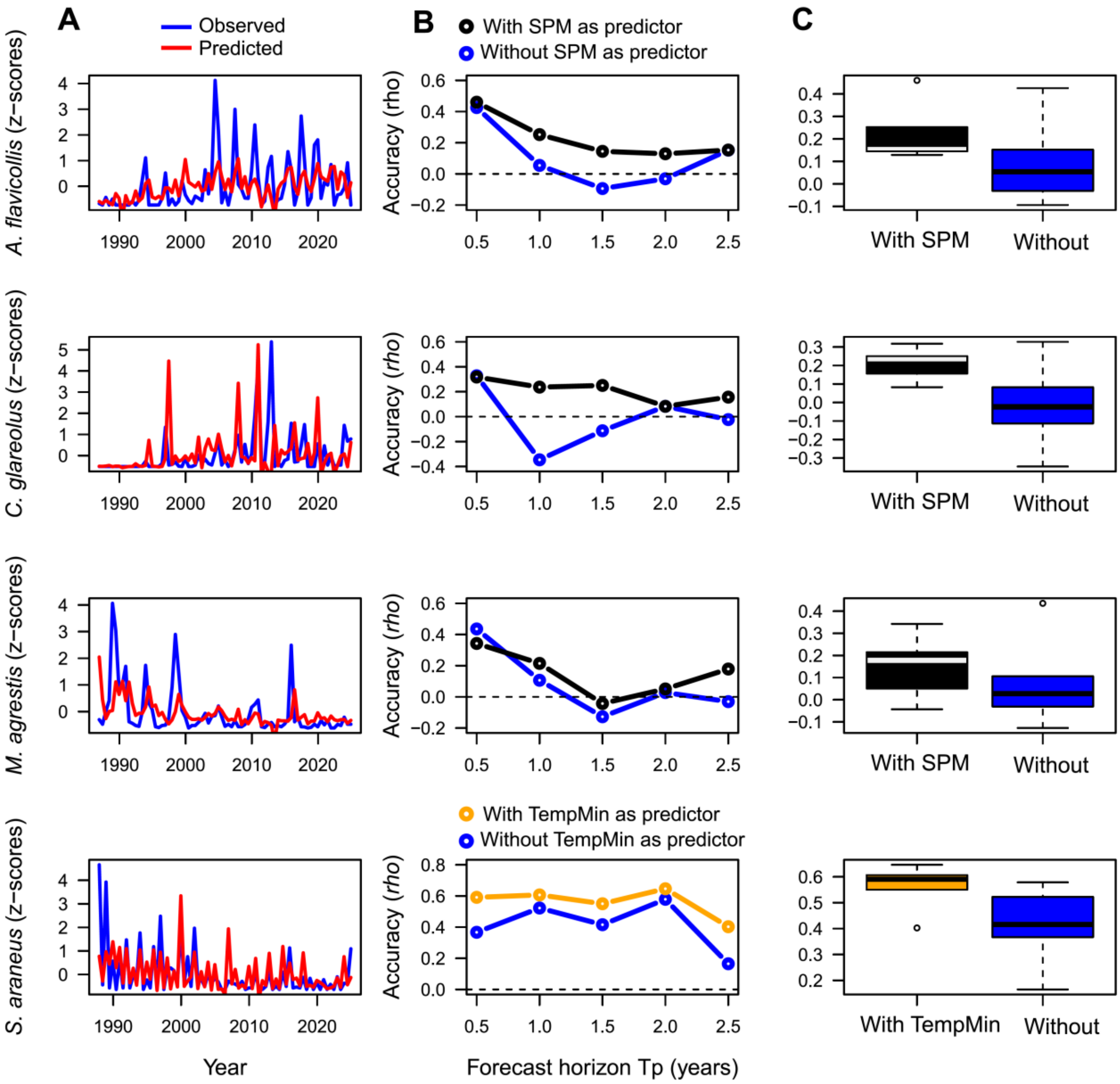


**Fig. S14.** Forecasting with or without key environmental drivers. (A) Comparison of observed vs predicted time series from the full models (leave-one-block-out- forecasts), in blue and red respectively at a forecast horizon of *Tp* = 1 year. We used multivariate S-maps with the abundance of the target species and its main competitors to forecast at horizons *Tp* 1 to 5 (i.e., 6 months to 2.5 years) as a baseline (blue) vs. with SPM levels (black) or min temperature (yellow) as additional predictors. (B) Forecast accuracy ($\rho$) as a function of forecast horizon *Tp* (in years), with (black, yellow) or without (blue) key drivers. (C) Comparisons of forecast accuracy with or without additional key drivers, using boxplots.

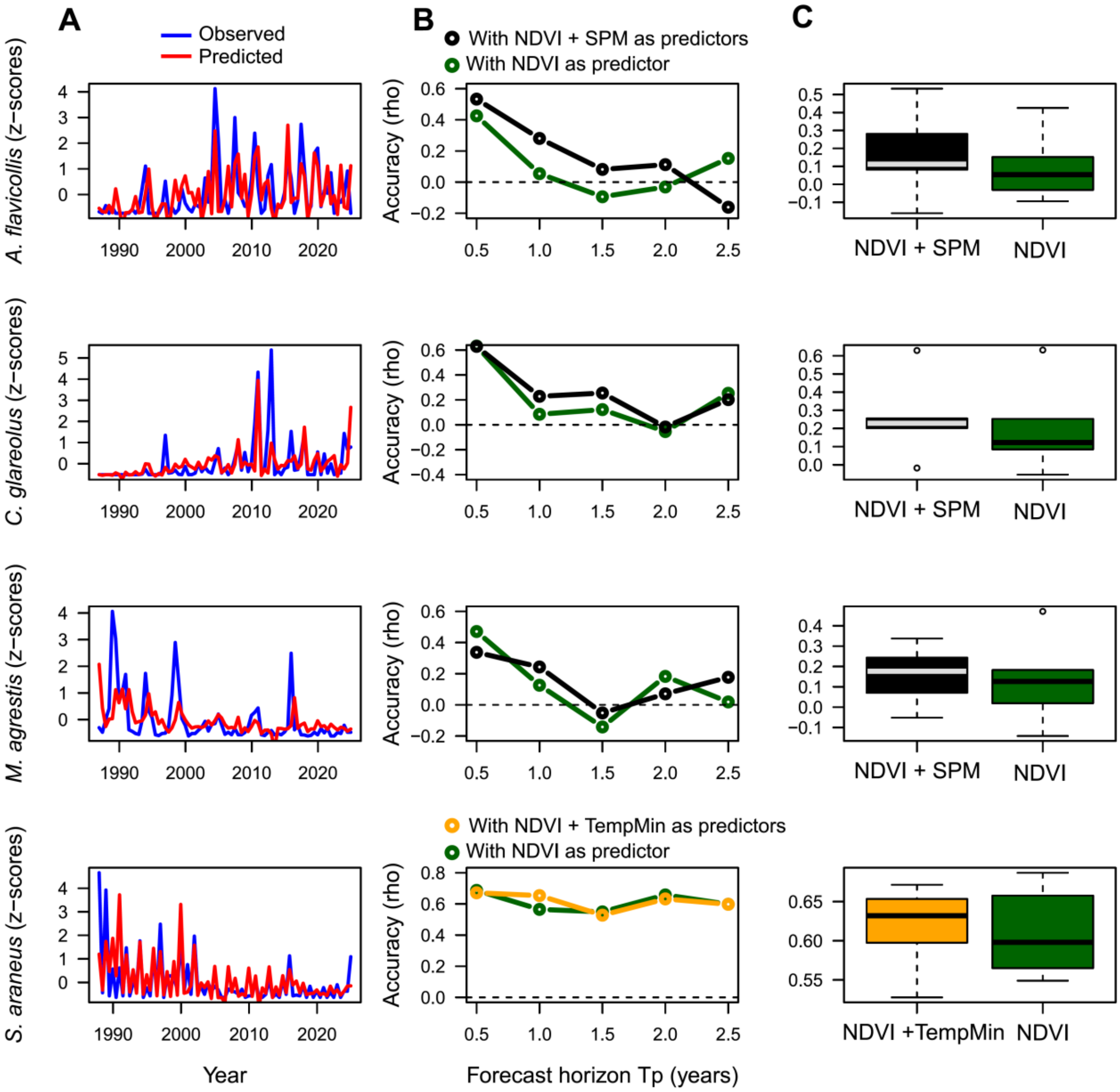


**Fig. S15**: (A) Comparison of observed vs predicted time series (leave-one-block-out- forecasts), in blue and red respectively. We used multivariate S-maps with the abundance of the target species and its main competitors to forecast at horizons *Tp* 1 to 5 (i.e., 6 months to 2.5 years), with an exclusion radius of 2, and NDVI as a baseline (dark green) vs. with these predictors and SPM levels. (B) Forecast accuracy (σ) as a function of forecast horizon *Tp* (in years), with (black) or without (green) SPM levels as predictor. Adding NDVI improves forecasts compared to no environmental predictors (forecasts in blue, Fig. S15, but adding SPM levels tend to improve forecasts (this figure). (C) Comparisons of forecast accuracy using boxplots.

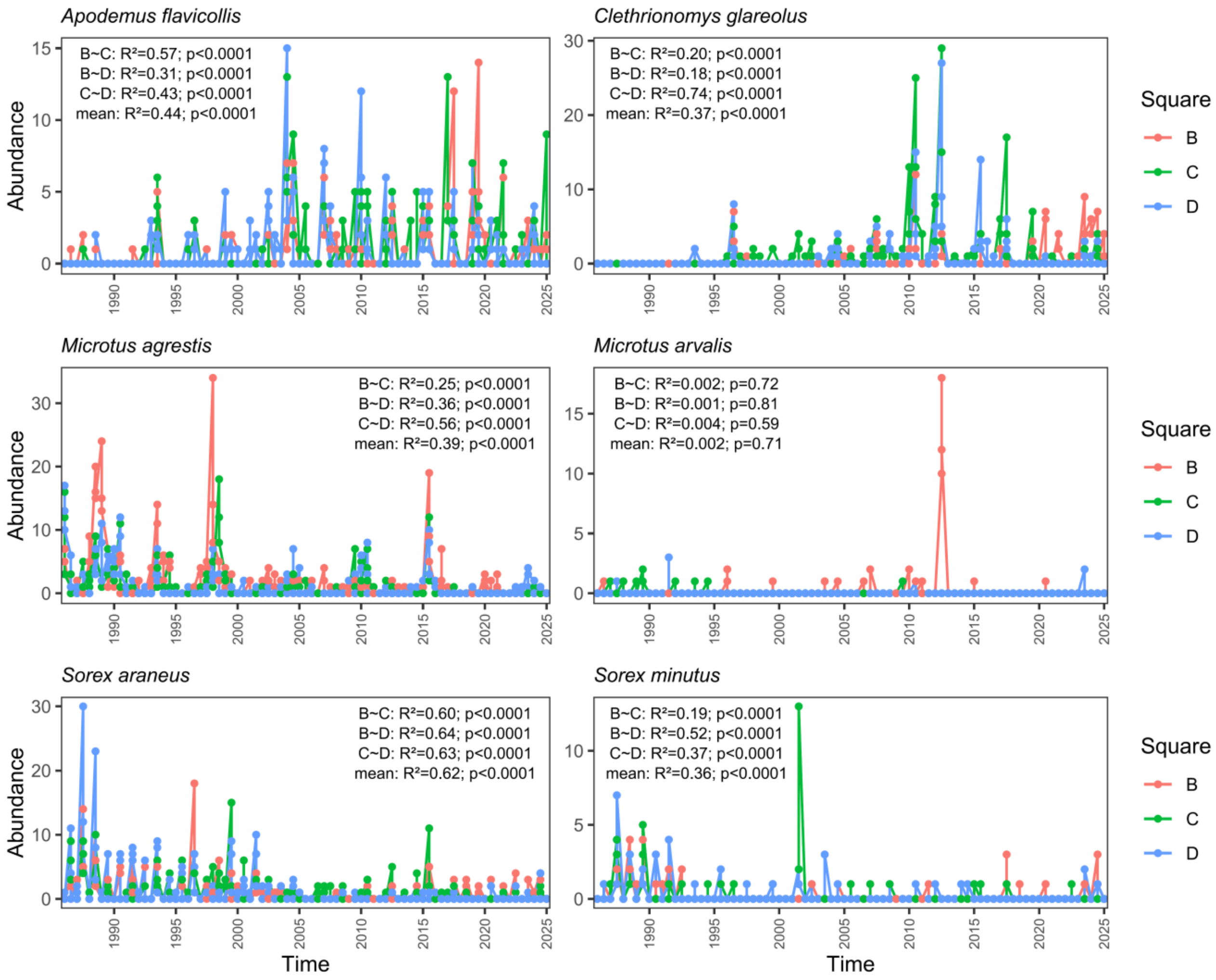


**Fig. S16:** Small- mammal time series displayed at the plot level ("Square" B, C, and D, in red, green, and blue, respectively). Statistics of relationships among time series within species and between pairs of plots (B–C, C–D, and B–D) are displayed as text insets ($R^2$ and p-values).

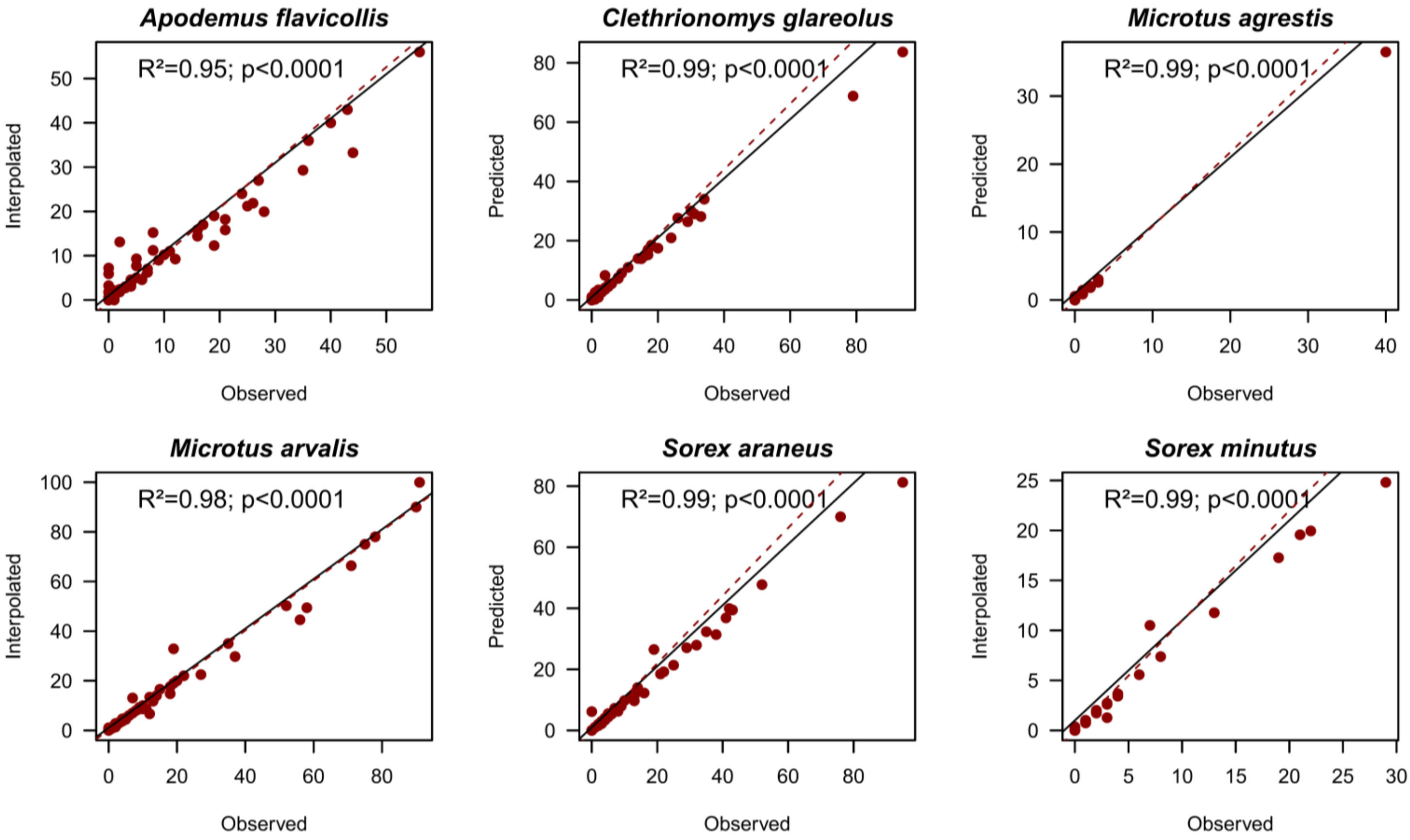


**Fig. S17:** Relationships between original (observed) and cubic Hermite (interpolated) time series (after pooling of each plot in Supplementary Fig. S17). Dashed red lines are the fit of the linear OLS regression, and the black line shows the proportional line (1:1). Statistics of relationships between observed and interpolated time series for each species are displayed as text insets ($R^2$ and p-values).

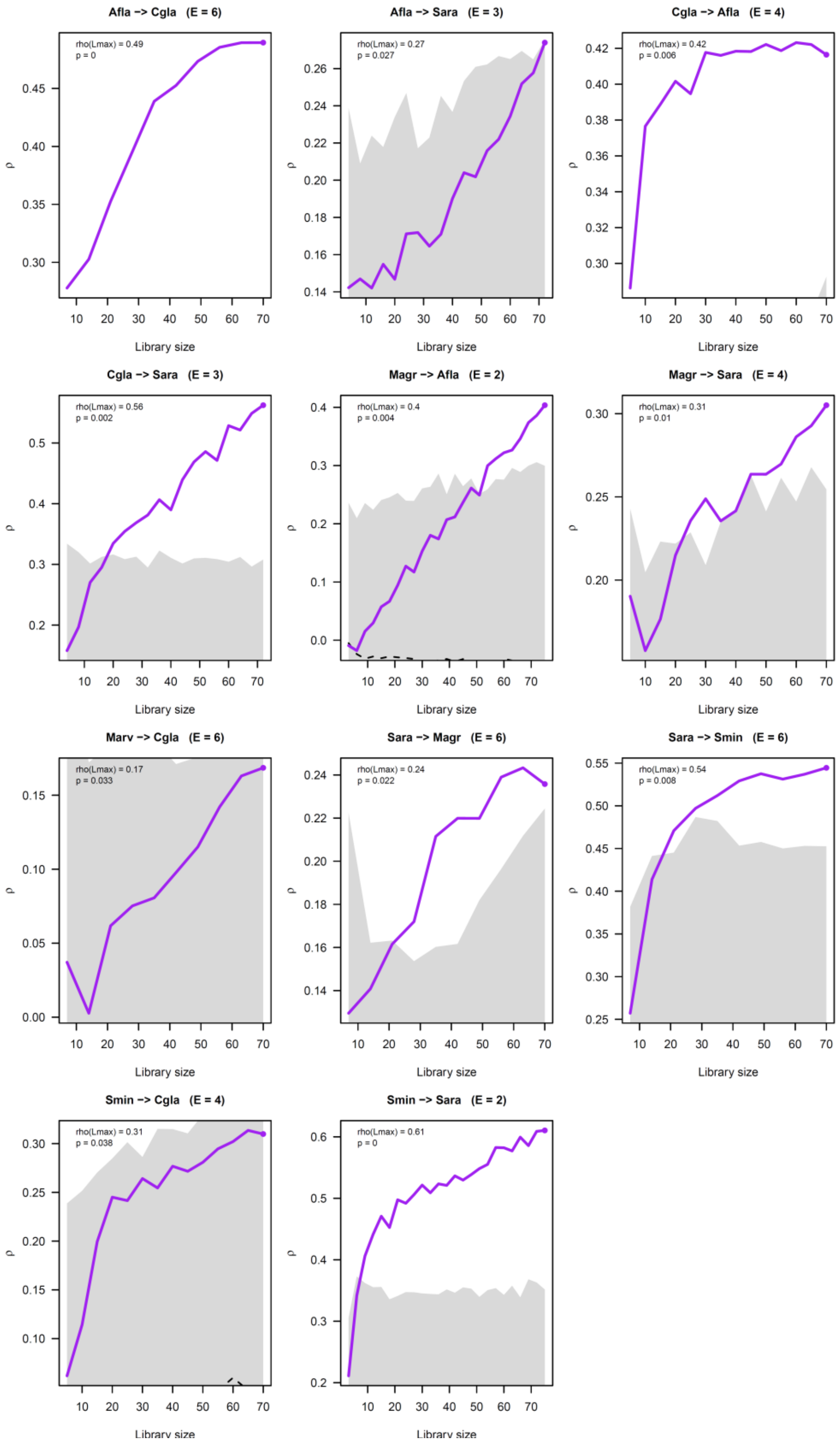


**Fig. S18**: Convergent-cross mapping (CCM) plots showing the convergence of observed prediction skills (purple line) as library size *L* increases for each of the retained significant pairs of species interactions in the simplified network presented in Fig. 2C of the main text. Grey bands show the 95% null distribution based on 1000 seasonal surrogates, and p-values indicate that observed skill at max *L* is superior to 95% of the skills drew from this null distribution. Embedding dimensions *E* are given in brackets.

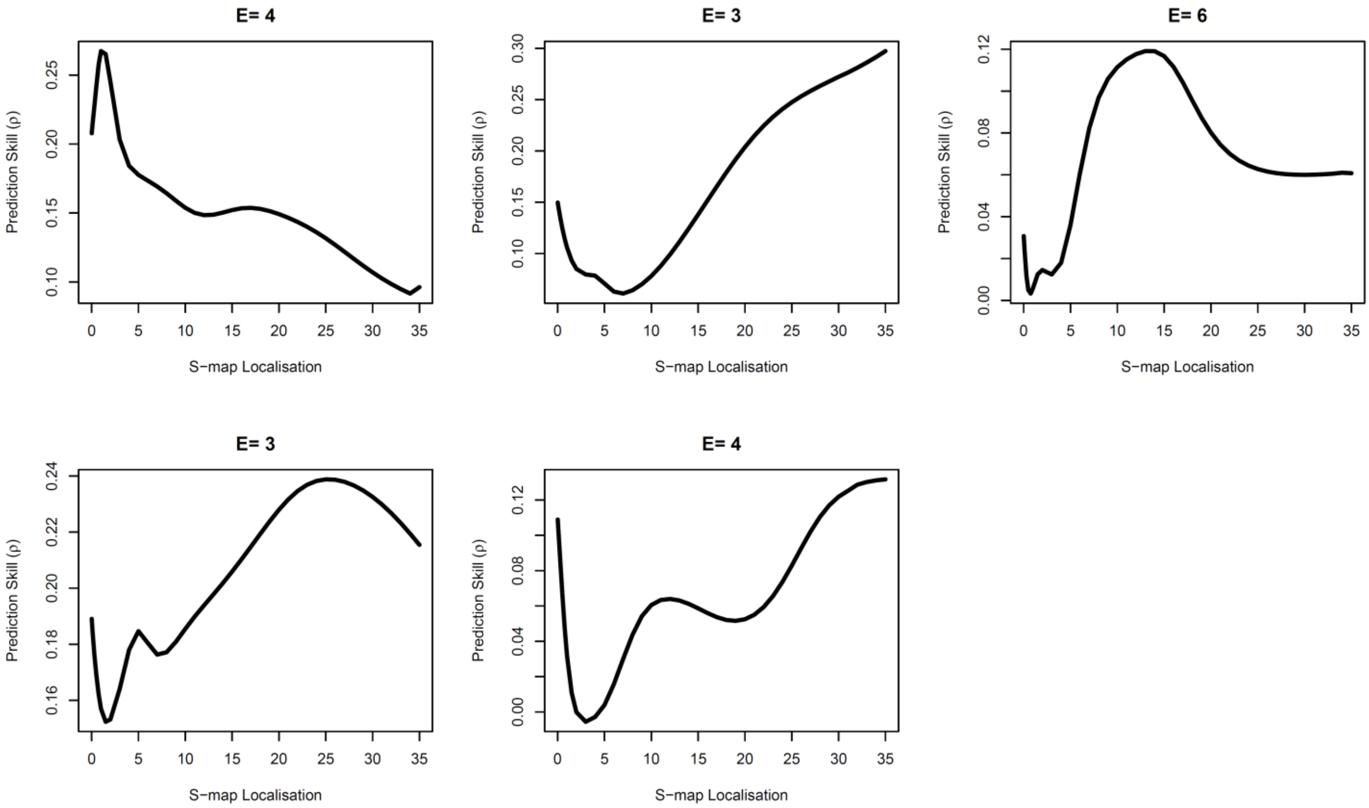


**Fig. S19**: S-map optimization plots of the prediction skill *ρ* as a function of the S-map localisation (or nonlinearity parameter *θ*) for the multivariate S-maps used to reconstruct the full matrix of time varying interaction strengths in the full network of 5 species (i.e., excluding *M. arvalis*; 20 pairs). Species are given from top left to bottom right: *A. flavicollis*, *C. glareolus*, *M. agrestis*, *S. araneus* and *S. minutus*. Clear non-linear multivariate state spaces are found for *A. fla.* (*θ = 0.30)*, *M. agr.* (*θ= 13)*, and *S. ara.* (*θ* = 25). For *C. glareolus* and *S. minutus*, we setted *θ* to 15 to avoid hyperlocal fitting. Embedding dimensions *E* are given on the top of each plot.

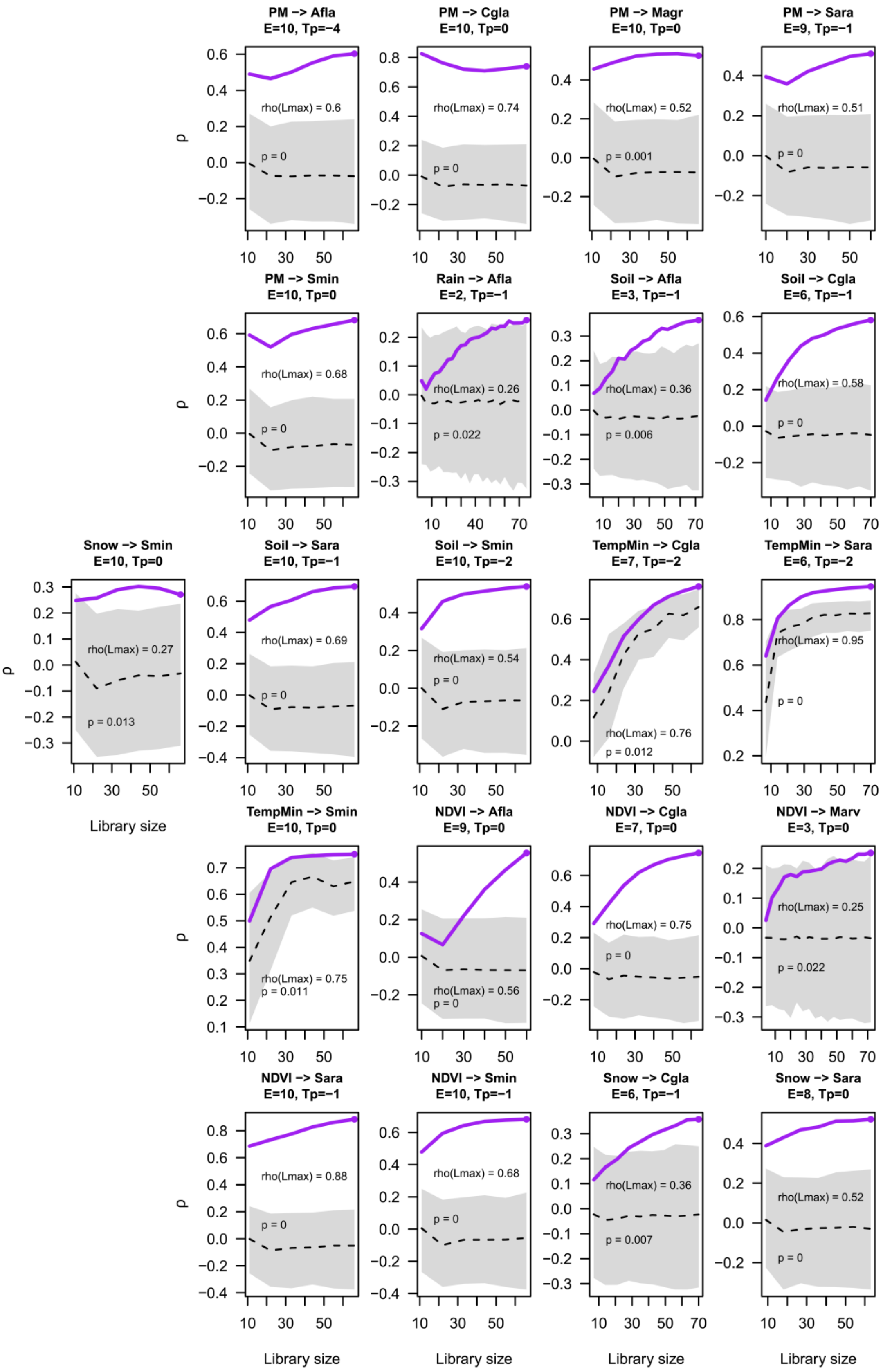


**Fig. S20**: Convergent-cross mapping (CCM) plots showing the convergence of observed prediction skills (purple line) as library size increases for each significant environmental driver → species causal link. Grey bands show the 95% null distribution based on 1000 seasonal surrogates.

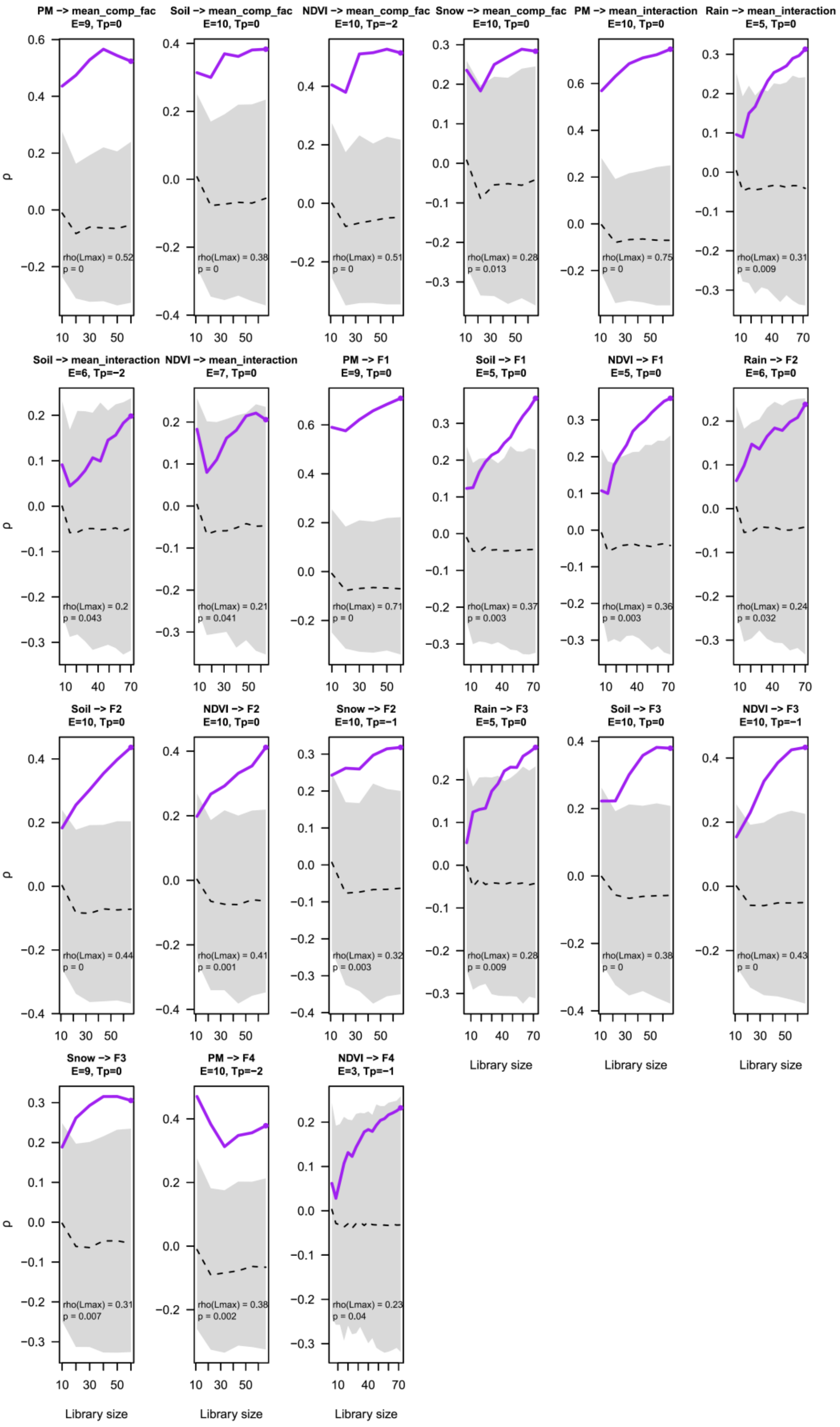


**Fig. S21**: Convergent-cross mapping (CCM) plots showing the convergence of observed prediction skills (purple line) as library size increases for each significant environmental driver → interaction metric causal link. Grey bands show the 95% null distribution based on 1000 seasonal surrogates. "PM" stands for SPM, "Soil" stands for soil moisture, "mean_comp_fac" stands for mean of signed interactions, and "mean_interactions" for mean of absolute interactions.

**Table S1**: Structure of the different multivariate S-maps used for the different aspects of the study: A) Interactions, B) Forecasting, C) Response surfaces, with the target (response) and column(s) (predictors), the library size *L*, the best embedding dimension *E*, the forecast horizon *Tp* (for "forecasting", we ran S-maps from *Tp* = 1 to 5), the nonlinearity parameter theta, and the predictive skill at *Tp* for forecasting are given as columns.

| Multivariate S-maps | Target | Column(s) | *L* | *E* | *Tp* | *θ* | *ρ* |
|---|---|---|---|---|---|---|---|
| ***A) Interactions*** | ***Afla*** | ***~ Afla*** | **1–78** | **1** | **2** | **1.5** | |
| (intraspecific) | ***Cgla*** | ***~ Cgla*** | **1–78** | **1** | **2** | **7** | |
| | ***Magr*** | ***~ Magr*** | **1–78** | **1** | **2** | **0.75** | |
| | ***Sara*** | ***~ Sara*** | **1–78** | **1** | **2** | **1.5** | |
| | ***Smin*** | ***~ Smin*** | **1–78** | **1** | **2** | **7** | |
| (interspecific) | ***Afla*** | ***~ Cgla + Magr + Sara + Smin*** | **1–78** | **4** | **0** | **1** | |
| | ***Cgla*** | ***~ Afla + Magr + Marv + Sara + Smin*** | **1–78** | **3** | **0** | **15** | |
| | ***Magr*** | ***~ Afla + Cgla + Sara + Smin*** | **1–78** | **6** | **0** | **13** | |
| | ***Sara*** | ***~ Magr + Afla + Cgla + Smin*** | **1–78** | **3** | **0** | **25** | |
| | ***Smin*** | ***~ Sara + Afla + Cgla + Magr*** | **1–78** | **4** | **0** | **15** | |
| ***B) Forecasting*** | ***z-Afla*** | ***~ z-Afla + z-Cgla*** | **1–78** | **5** | **1** | **0.75** | **0.4** |
| (exclusion radius: 2) | ***z-Afla*** | ***~ z-Afla + z-Cgla + z-SPM*** | **1–78** | **1** | **1** | **1** | **0.5** |
| | ***z-Cgla*** | ***~ z-Cgla + z-Afla*** | **1–78** | **6** | **1** | **0.1** | **0.3** |
| | ***z-Cgla*** | ***~ z-Cgla + z-Afla + z-SPM*** | **1–78** | **6** | **1** | **8** | **0.3** |
| | ***z-Magr*** | ***~ z-Magr + z-Afla + z-Cgla*** | **1–78** | **4** | **1** | **0.75** | **0.4** |
| | ***z-Magr*** | ***~ z-Magr + z-Afla + z-Cgla + z-SPM*** | **1–78** | **6** | **1** | **0.1** | **0.3** |
| | ***z-Sara*** | ***~ z-Sara_2 + z-Afla_2*** | **1–78** | **5** | **1** | **1.5** | **0.4** |
| | ***z-Sara*** | ***~ z-Sara_2 + z-Afla_2 + z-MinTemp*** | **1–78** | **5** | **1** | **3** | **0.6** |
| | ***z-Afla*** | ***~ z-Afla + z-Cgla + z-NDVI*** | **1–78** | **4** | **1** | **1.5** | **0.4** |
| | ***z-Afla*** | ***~ z-Afla + z-Cgla + z-NDVI + z-SPM*** | **1–78** | **4** | **1** | **6** | **0.5** |
| | ***z-Cgla*** | ***~ z-Cgla + z-Afla + z-NDVI*** | **1–78** | **2** | **1** | **3** | **0.6** |
| | ***z-Cgla*** | ***~ z-Cgla + z-Afla + z-NDVI + z-SPM*** | **1–78** | **2** | **1** | **3** | **0.6** |
| | ***z-Magr*** | ***~ z-Magr + z-Afla + z-Cgla + z-NDVI*** | **1–78** | **1** | **1** | **1.5** | **0.5** |
| | ***z-Magr*** | ***~ z-Magr + z-Afla + z-Cgla + z-NDVI + z-SPM*** | **1–78** | **3** | **1** | **0.01** | **0.3** |
| | ***z-Sara*** | ***~ z-Sara_2 + z-Afla_2 + z-NDVI*** | **1–78** | **2** | **1** | **3** | **0.7** |
| | ***z-Sara*** | ***~ z-Sara_2 + z-Afla_2 + z-NDVI + z-MinTemp*** | **1–78** | **2** | **1** | **3** | **0.7** |
| ***C) Response surfaces*** | **F1** | **~ SPM + NDVI** | **1–78** | **1** | **0** | **0.3** | |
| (exclusion radius: 12) | **F2** | **~ Soil + NDVI** | **1–78** | **9** | **0** | **0.5** | |
| | **F3** | **~ Soil + NDVI** | **1–78** | **13** | **0** | **0.1** | |
| | **F4** | **~ SPM + NDVI** | **1–78** | **6** | **0** | **0.01** | |
| | **Abs. Int. Strength** | **~ SPM + NDVI** | **1–78** | **10** | **0** | **0.01** | |
| | **Mean Interactions** | **~ SPM + NDVI** | **1–78** | **1** | **0** | **0.01** | |
| | **LLE** | **~ Soil + F2** | **1–67** | **15** | **0** | **0.01** | |
| | **LLE** | **~ NDVI + F3** | **1–67** | **13** | **0** | **1.5** | |
| | **LLE** | **~ SPM + Abs_Inter.** | **1–67** | **4** | **0** | **0.75** | |
